# Cell non-autonomous control of autophagy and metabolism by glial cells

**DOI:** 10.1101/2022.12.15.520639

**Authors:** Melissa G. Metcalf, Samira Monshietehadi, Arushi Sahay, Ashley E. Frakes, Jenni Durieux, Martina Velichkovska, Cesar Mena, Amelia Farinas, Melissa Sanchez, Andrew Dillin

## Abstract

Glia are the protectors of the nervous system, providing neurons with support and protection from cytotoxic insults. We previously discovered that four astrocyte-like glia can regulate organismal proteostasis and longevity in *C. elegans*. Expression of the UPR^ER^ transcription factor, XBP-1s, in these glia increases stress resistance, longevity, and activates the UPR^ER^ in intestinal cells via neuropeptides. Autophagy, a key regulator of metabolism and aging, has been described as a cell autonomous process. Surprisingly, we find that glial XBP-1s enhances proteostasis and longevity by cell non-autonomously reprogramming organismal lipid metabolism and activating autophagy. Glial XBP-1s regulates the activation of another transcription factor, HLH-30/TFEB, in the intestine. HLH-30 activates intestinal autophagy, increases intestinal lipid catabolism, and upregulates a robust transcriptional program. Our study reveals a novel role for glia in regulating peripheral lipid metabolism, autophagy, and organellar health through peripheral activation of HLH-30 and autophagy.

## INTRODUCTION

The central nervous system senses and processes internal and afferent signals which convey information about nutrient status and stress within an organism. The ability of the brain to accurately detect and correctly respond to these signals is essential for the survival of all animals. Both glial cells and glia-neuronal contact sites are hubs for integrating metabolic cues (Garcia-Caceres et al., 2019). Signals sent from the nervous system, after detecting changes in internal or external environments, have been shown to modulate lifespan, regulate protein homeostasis (proteostasis), and stress resistance (Brandt et al., 2018; Durieux et al., 2011; Frakes et al., 2020; Taylor and Dillin, 2013; Williams et al., 2014). As an organism ages, the ability of a cell to detect and respond to cellular and organismal stressors declines. Thus, the ability of the nervous system to detect and respond to signals from the environment and inter-organ communication can shape the progression of the aging process and the onset of age associated diseases (Camandola and Mattson, 2017; Mayorga-Weber et al., 2022).

A cell employs numerous mechanisms to respond to cellular insults and maintain proteostasis through the coordinated action of distinct stress response pathways in their organelles and cellular compartments (Hipp et al., 2019). A key stress response pathway is found in the endoplasmic reticulum (ER), an organelle which has a central role in lipid and protein biosynthesis, being the site of production of all the transmembrane proteins and lipids for the majority of a cell’s organelles (Oakes and Papa, 2015). Increased protein or lipid demands within a cell disrupt the function of the ER and increase ER stress, triggering the unfolded protein response of the ER (UPR^ER^) to restore cellular homeostasis (Metcalf et al., 2020). The IRE-1 branch of the UPR^ER^ detects unfolded proteins in the ER lumen and lipid perturbations in the ER membrane, leading to the regulated splicing of the mRNA encoding the transcription factor XBP-1 (Calfon et al., 2002; Cox et al., 1997; Fu et al., 2012; Kono et al., 2017; Promlek et al., 2011; Sidrauski and Walter, 1997). Spliced XBP-1, XBP-1s, is subsequently translated, leading to the regulation of transcriptional targets to reduce ER stress by repressing protein translation, regulating lipid synthesis, inducing protein chaperone expression, and eliminating misfolded and damaging proteins (Adamson et al., 2016; Piperi et al., 2016; Yamamoto et al., 2007).

UPR^ER^ activation in the nervous system can be non-autonomously communicated to other tissues to regulate organismal stress resistance and longevity (Taylor and Dillin, 2013; Williams *et al*., 2014). Pan-neuronal activation of the UPR^ER^ leads to organismal effects on lysosomal and lipid metabolism via neurotransmitter signaling in the roundworm *Caenorhabditis elegans* (Daniele et al., 2020; Higuchi-Sanabria et al., 2020; Imanikia et al., 2019a; Imanikia et al., 2019b; Ozbey et al., 2020). *C. elegans* has a simple nervous system, with 302 neurons and 56 glial cells encompassing close to one third of total cells in the organism (Singhvi and Shaham, 2019). Of the 56 glial cells, the four glial cells of the cephalic sheath (CEPsh) are positioned around the brain of *C. elegans*, the ‘nerve ring’, with their anterior processes forming a sensory organ at the mouth of the organism (Singhvi and Shaham, 2019). CEPsh glia have numerous functions that parallel mammalian glial cell types, including regulating the interface between peripheral organs and the nervous system, providing trophic support, and acting as neuronal and glial progenitors (Katz et al., 2018; Katz et al., 2019; Lago-Baldaia et al., 2020; Rapti et al., 2017; Singhvi and Shaham, 2019). Over-expression of a UPR^ER^ transcription factor, XBP-1s, in CEPsh glia leads to lifespan extension and ER stress resistance through induction of the UPR^ER^ in distal intestinal cells through release of dense core vesicles (DCV) and neuropeptides (Frakes *et al*., 2020). These findings indicate that this subset of glial cells can regulate systemic proteostasis, interorgan communication, and organismal aging through neuropeptide signaling. However, the downstream mechanisms regulating these phenotypes remain unknown.

Here, we find that ectopic expression of constitutively active XBP-1s in four CEPsh glial cells initiates a robust metabolic program, leading to lipid depletion and ER remodeling peripherally. Cell non-autonomous signaling from CEPsh glia activates intestinal HLH-30, a transcription factor important for lysosomal and autophagy homeostasis (Lapierre et al., 2013). Interestingly, HLH-30 is required for the lipid depletion, lifespan extension, and increased proteostasis of glial XBP-1s animals. In parallel to HLH-30 activation in the periphery, glial XBP-1s animals have activated autophagy in intestinal tissue. Autophagy is required for the extension of longevity and increased ER remodeling in the intestine of glial XBP-1s animals. Finally, glial XBP-1s signaling to activate HLH-30 in the periphery requires dense core vesicle release. Taken together, we find that cell non-autonomous signaling from XBP-1s expressing CEPsh glia activates a peripheral metabolic program required for the beneficial impacts on organismal proteostasis, organellar health, and longevity.

## RESULTS

### Glial XBP-1s animals have reduced lipid content, droplet number, and increased lysosomes in their periphery

The UPR^ER^ and XBP-1 have been shown to regulate lipid metabolism, and changes in lipid metabolism regulate lifespan. (Lopez-Otin et al., 2016; Singh et al., 2019; Templeman and Murphy, 2018). To determine if the long-lived and stress resistant animals overexpressing the coding sequence of *xbp-1s* in CEPsh glia (glial XBP-1s) altered peripheral lipid metabolism, we measured whole-body total neutral lipid content via BODIPY 493/503 staining. We found that glial XBP-1s animals had significantly reduced staining of total neutral lipids compared to wild-type animals, suggesting that glial UPR^ER^ activation leads to reduced peripheral lipid stores (Figure 1A and 1B), which is not due to differences in background autofluorescence (Figure S1A). When comparing neutral lipid content to animals with known fat storage defects, *eat-2(e1372)* and *daf-7(ad1116)*, show increased and decreased BODIPY 493/503, respectively (Figure S1B) (Klapper et al., 2011).

**Figure 1.**
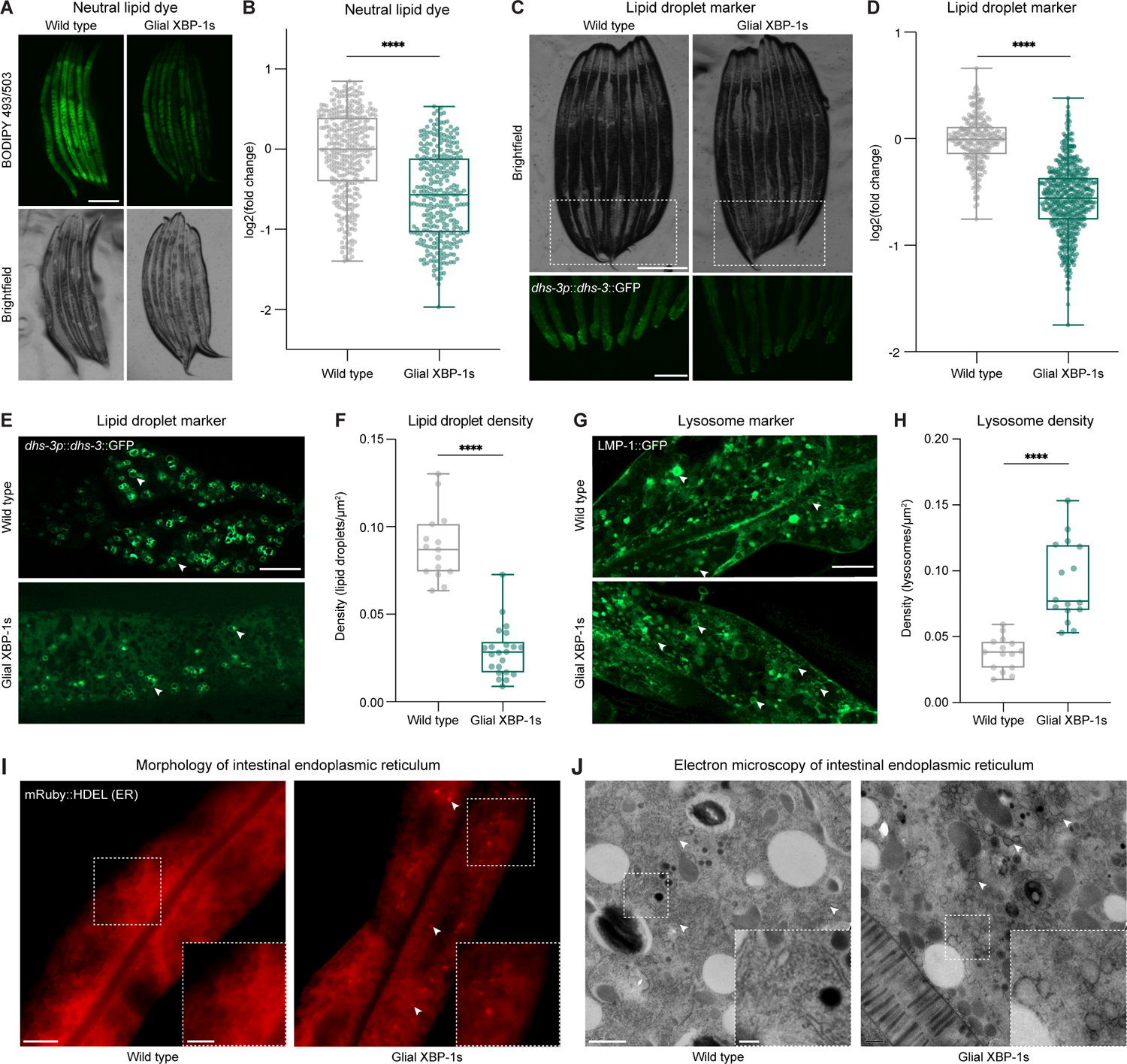
Expression of XBP-1s in CEPsh glia modulates peripheral lipid metabolism and ER remodeling in *C. elegans*. (A) Fluorescent light micrographs of wild-type (N2) and glial XBP-1s animals stained with BODIPY 493/503 dye and imaged at day 2 of adulthood. Scale bar, 250 µm. (B) Whole animal fluorescence intensity quantification of BODIPY 493/503 dye in day 2 adults in wild-type (grey) and glial XBP-1s animals (green) using COPAS BioSorter. p <0.0001 (****), using non-parametric Mann-Whitney test. Plots are representative of three biological replicates, n = 362 (wild type) and n = 302 (glial XBP-1s). Results were normalized to mean fluorescent intensity of wild-type animals. Each dot represents one animal, and the boxplot shows median (horizontal line), the first and third quartiles (box), and smallest and largest data points (whiskers). (C) Fluorescent light micrographs of wild-type and glial XBP-1s animals of intestinal lipid droplets (*dhs-3*p::*dhs-3*::GFP) imaged at day 2 of adulthood. Scale bar, 250 µm. (D) Whole animal fluorescence intensity quantification of intestinal *dhs-3*p::*dhs-3*::GFP lipid droplet marker in wild-type (grey) and glial XBP-1s animals (green) using COPAS BioSorter. p < 0.0001 (****), using non-parametric Mann-Whitney test. Plots are representative of three biological replicates, n = 254 (wildtype) and n = 532 (glial XBP-1s). Results were normalized to mean fluorescent intensity of wild-type animals transgenic for *dhs-3*p::*dhs-3*::GFP. (E) Representative Airyscan micrographs of *dhs-3*p::*dhs-3*::GFP labeled lipid droplets from anterior intestines of day 2 wild-type and glial XBP-1s adult animals. Arrowheads point at lipid droplets. Scale bar, 10 µm. (F) Quantification of intestinal *dhs-3*p::*dhs-3*::GFP labeled lipid droplets in wild-type (grey) and glial XBP-1s (green) animals. Lipid droplets were counted from three independent replicates using ImageJ and expressed as density. Density was determined by dividing the number of lipid droplets by the total area expressing *dhs-3*p::*dhs-3*::GFP in each respective image. N = 15 (wild type) and N = 22 (glial XBP-1s). p < 0.0001 (****), using non-parametric Mann-Whitney test. (G) Representative Airyscan micrographs of *lmp-1*::GFP labeled lysosomes in the anterior intestine of day 2 adults in both wild-type and glial XBP-1s animals. Arrowheads point at lysosomes. Scale bar, 10 µm. (H) Quantification of intestinal *lmp-1*::GFP labeled lysosomes in wild-type (grey) and glial XBP-1s (green) animals. Lysosomes were counted from three independent replicates using ImageJ and expressed as density. Density was determined by dividing the number of lysosomes by the total area that expressed *lmp-1*::GFP in each respective image. N = 15 (wild type) and N = 16 (glial XBP-1s). p < 0.0001 (****), using non-parametric Mann-Whitney test. (I) Representative confocal micrographs of intestinal ER morphology (*vha-6p*::ERss::mRuby::HDEL, ERss = *hsp-4* ER signal sequence), wild-type and glial XBP-1s animals were imaged at day 2 of adulthood. Arrowheads mark ER puncta. Scale bar, 10 µm. Scale bar of inset, 5 µm. (J) Electron micrographs of intestine from wild-type and glial XBP-1s animals at day 2 of. Imaging was replicated in triplicate, age matched at Day 2, for a total of 15-20 animals being imaged per condition. Arrowheads mark rough endoplasmic reticulum. Scale bar, 1µm. Scale bar of inset, 0.2 µm.

Lipid droplets are a conserved organelle for fat storage and the main storage site of intestinal lipids in *C. elegans*. We hypothesized that lipid droplet content may be reduced in glial XBP-1s animals (Mak, 2012). We visualized lipid droplets using an intestinally-expressed green fluorescent protein (GFP)-tagged construct of a short-chain dehydrogenase, DHS-3, which localizes to lipid droplet membranes in *C. elegans* (Na et al., 2015). Using this marker, we found that glial XBP-1s animals had reduced fluorescence intensity of the DHS-3::GFP lipid droplet marker when compared to wild-type animals (Figure 1C and 1D), and that this decrease was not due to differences in basal autofluorescence (Figure S1C). Further, we observed a decrease in lipid droplet density, determined by the number of lipid droplets per area of intestine imaged, in the intestines of glial XBP-1s animals compared to wild-type animals (Figure 1E and 1F). These data point to the intestine of glial XBP-1s animals, the same organ which receives a signal to activate the UPR^ER^, having reduced lipid content.

We hypothesized that the reduced lipid content in glial XBP-1s animals may be due to increased lipid catabolism by lysosomes. Lysosomes act as cellular regulators of energy metabolism, functioning as both catabolic recycling centers within a cell and metabolic signaling and sensing hubs that govern cell growth decisions (Lawrence and Zoncu, 2019; Thelen and Zoncu, 2017). We investigated if there was an increase in lysosome content in the intestine of glial XBP-1s animals, which may be indicative of increased lipid catabolism (Singh et al., 2009). Lysosome density was measured using a GFP-fusion protein of the *C. elegans* homolog of the lysosomal surface protein LAMP1, *lmp-1* (Figure 1G). Using this marker, we found that glial XBP-1s animals had significantly higher lysosome density in their intestine when compared to wild-type animals (Figure 1H). To confirm that the decrease in lipid content and increase in lysosome numbers was not due to a decrease in food intake, we compared feeding rates of glial XPB-1s animals compared to control. Glial XBP-1s animals have similar pumping rates to wild-type animals, indicating that they are consuming comparable amounts of their bacterial food source (Figure S1D) (Song and Avery, 2013). This increase in lysosomal number within the intestine of glial XBP-1s animals is indicative of an increase in organismal lipid catabolic and proteostatic activity.

### Glial XBP-1s animals have altered ER morphology and function in distal tissues

As the central hub for membrane biogenesis and the production of the majority of membrane lipids, the ER is particularly sensitive to changes in lipid content, which can drastically change its structure and function (Bernales et al., 2006; van Meer et al., 2008). Activation of the UPR^ER^ increases both the size of the ER and its folding capacity to reestablish homeostasis within the organelle (Bernales *et al*., 2006; Schuck et al., 2009; Sriburi et al., 2007; Sriburi et al., 2004). To test if glial UPR^ER^ activation affected peripheral ER morphology, we visualized the intestinal ER using a fusion protein, *vha-6p*::ERss::mRuby::HDEL, that localizes to the ER lumen through a heat shock protein 4 (HSP-4) signal sequence, is retained in the ER through a C-terminal HDEL sequence, and expresses a red fluorescent protein, mRuby. Interestingly, glial XBP-1s animals form puncta of mRuby::HDEL labelled ER that are not seen in wild-type animals (Figure 1I). Additionally, we found that glial XBP-1s animals had an increase in intestinal secretion of secretory proteins, visualized using VIT-2::GFP (Figure S1E). VIT-2::GFP is a fluorescent fusion protein of vitellogenin, a yolk protein that is produced in the intestinal ER of *C. elegans*, secreted and then endocytosed by developing eggs (Grant and Hirsh, 1999). The level of VIT-2::GFP fluorescence in egg cells is used as a proxy for ER function in the intestine of *C. elegans* (Daniele *et al*., 2020). To better understand the nature of these changes in the intestinal ER of glial XBP-1s animals, we performed high-magnification transmission electron microscopy (Figure 1J). We identified changes in rough ER morphology in glial XBP-1s animals, where rough ER formed circular-like structures compared to the reticulated ER structure in wild-type animals (Figure S1F). Taken together, these results demonstrate that glial XBP-1s animals have changes in ER morphology within their intestines, indicating expansion of the ER, as well as increased secretion of secretory proteins.

### HLH-30/TFEB is required for glial XBP-1s lifespan extension

The changes in storage fat levels, lipid droplet content, and increased lysosome number suggest that a master regulator of metabolism is activated in the periphery of these glial XBP-1s animals. Because these metabolic phenotypes are predictors of longevity of the glial XBP-1s animals, we asked which, if any, of the known master regulators of metabolic function might be required for the glial XBP-1s animals’ increased longevity. We found that the lifespan extension of glial XBP-1s animals was not dependent on key regulators of metabolism: *daf-16* (FOXO), *aak-1* and *aak-2* (AMPK), or *pha-4* (FOXA) (Figure 2A-D) (Apfeld et al., 2004; Greer et al., 2007; Lin et al., 1997; Ogg et al., 1997; Panowski et al., 2007). Surprisingly, glial XBP-1s animals had significantly upregulated levels of fluorescently tagged DAF-16 protein, DAF-16::GFP; however, nuclear localization of this transcription factor was not observed, suggesting that DAF-16 is not functioning to increase its target genes’ transcription (Figure S2A-B). HLH-30, the *C. elegans* ortholog of the mammalian transcription factor EB (TFEB), a master regulator of lysosomal biogenesis, autophagy, and lipid catabolism, was found to be required for glial XBP-1s lifespan extension (Lapierre *et al*., 2013; O’Rourke and Ruvkun, 2013; Settembre et al., 2013; Settembre et al., 2011). Knockdown of *hlh-30* via RNAi treatment suppressed the lifespan extension in glial XBP-1s animals (Figure 2E). In parallel, we found that loss of *hlh-30*, via a loss-of-function mutation *hlh-30(tm1978)*, suppressed the lifespan extension of glial XBP-1s animals (Figure 2F). These data point to a role for the transcription factor HLH-30 in mediating longevity in glial XBP-1s animals, and not other known regulators of metabolism.

**Figure 2.**
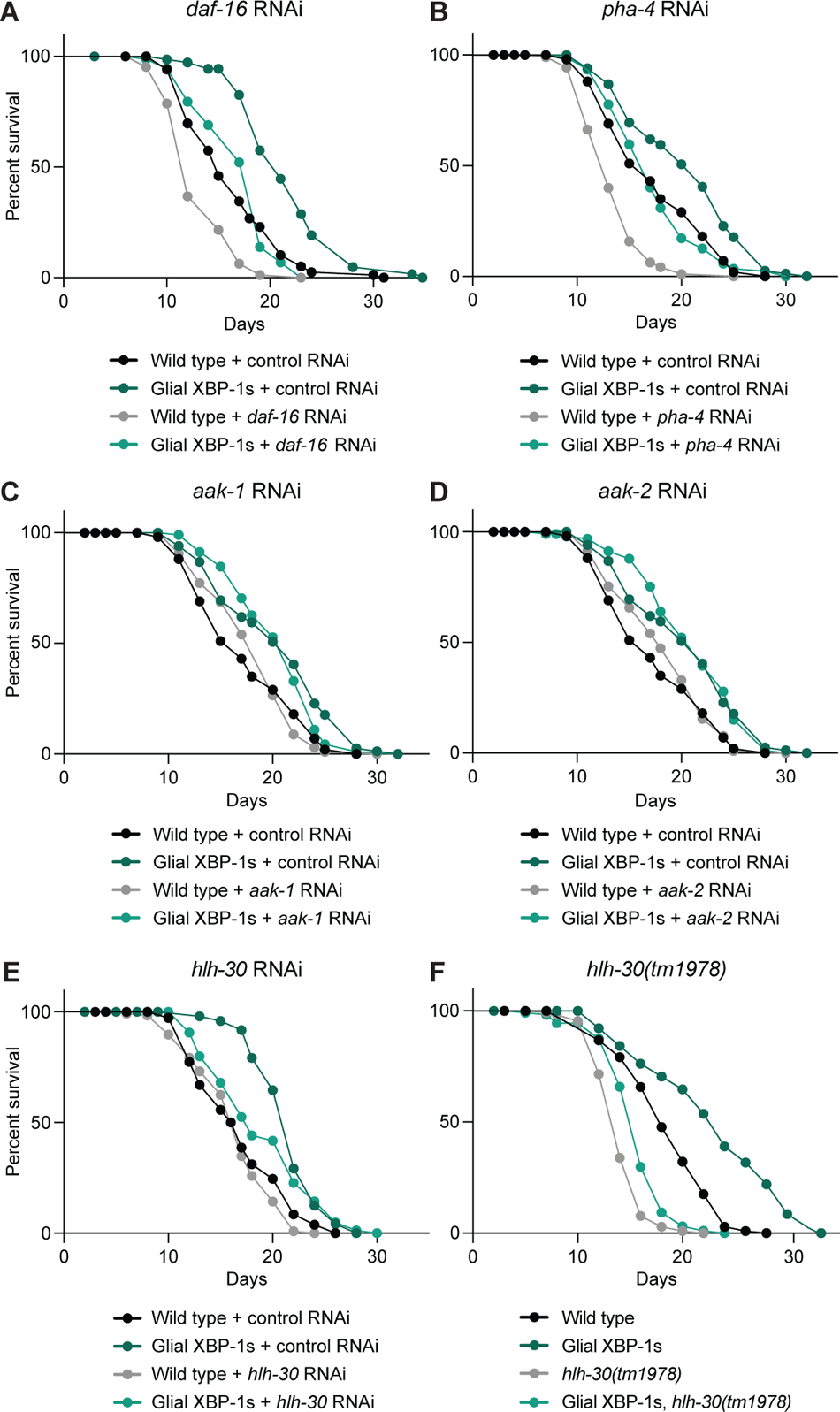
Lifespan screen of key metabolic regulators reveals HLH-30 requirement for longevity of glial XBP-1s animals. (A-E) Survival of wild-type (N2) and glial XBP-1s animals on control (HT115 *E. coli* expressing empty vector), *daf-16* (A), *aak-1* (B), *aak-2* (C), *pha-4* (D), and *hlh-30* (E) RNAi from L4 at 20°C. Lifespan is representative of three replicates. Graphs were plotted as Kaplan-Meier survival curves, and p values were calculated by Mantel-Cox log-rank test. See Table S1 for lifespan statistics. (F) Survival of wild-type and glial XBP-1s animals on control RNAi at 20°C, with and without *hlh-30(tm1978)* mutation. Graph was plotted as Kaplan-Meier survival curves, and p values were calculated by Mantel-Cox log-rank test. See Table S1 for lifespan statistics.

### HLH-30/TFEB is activated cell non-autonomously in peripheral tissue of glial XBP-1s animals via DCV release

HLH-30 is activated under times of stress to meet the metabolic needs of an organism with the cellular resources available. When HLH-30 is inactive, it is sequestered in the cytosol via scaffolding proteins, and upon activation via phosphorylation, HLH-30 moves from the cytoplasm into the nucleus. We used a fluorescent fusion protein, HLH-30::GFP, to determine levels of nuclear localization of HLH-30::GFP in the intestine of glial XBP-1s animals as a proxy for HLH-30 activation (Lapierre *et al*., 2013). We observed HLH-30::GFP to be enriched prominently in the nuclei of intestinal cells in adult glial XBP-1s animals (Figure 3A – 3C). These data indicate that HLH-30 is activated peripherally in the intestine when *xbp-1s* is constitutively active in CEPsh glia.

**Figure 3.**
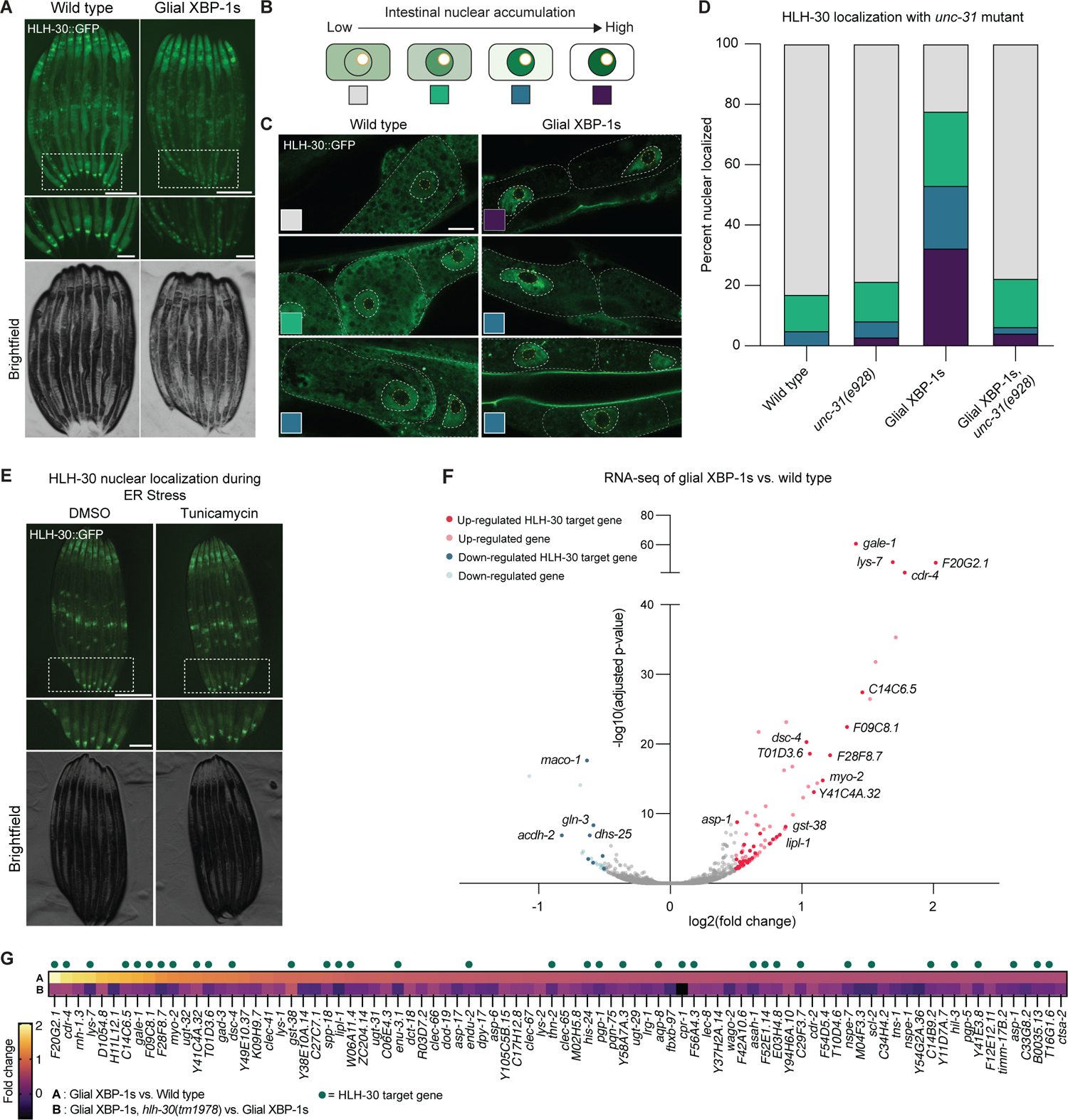
Glial XBP-1s cell non-autonomously activates intestinal HLH-30 which transcriptionally regulates genes involved in lipid catabolism and lysosomal biogenesis. (A) Fluorescent light micrographs of wild-type and glial XBP-1s animals transgenic for fluorescently tagged HLH-30 (HLH-30::GFP) grown on OP50 and imaged at day 2 of adulthood. HLH-30 translocates to the nucleus in glial XBP-1s animals. Inset shows posterior intestine. Scale bar, 250µm. Scale bar of inset, 100µm. (B) Schematic showing how HLH-30::GFP nuclear localization was scored. Categories include no nuclear enrichment (gray), weak nuclear enrichment (green), medium nuclear enrichment (blue), and strong nuclear enrichment (purple). Yellow circle indicates nucleolus, which transcription factors are excluded from. (C) Representative Airyscan micrographs of posterior intestinal cells of wild-type and glial XBP-1s animals transgenic for HLH-30::GFP grown on OP50 and imaged at day 2 of adulthood. Colored inset squares are representative of the scoring categories from (B). Scale bar, 10µm. Intestinal cells are outlined with a dashed grey line, intestinal nuclei are outlined with a dashed white line, and intestinal nucleoli are outlined with a dashed yellow line. (D) Nuclear translocation of HLH-30::GFP in anterior intestinal cells of day 2 adult wild-type and glial XBP-1s animals with and without the loss-of-function *unc-31* mutant, *unc-31(e928)*, N > 40 animals per condition from three independent replicates. Statistics done by Chi-squared test for independence with adjusted residual and Bonferroni correction, statistics shown in Table S5. (E) Representative fluorescent micrograph of wild-type animals transgenic for HLH-30::GFP treated either DMSO or tunicamycin. Scale bar, 250 µM. Scale bar of inset, 100 µM. Image is representative of two independent replicates, in n > 20 no animals showed nuclear localization. (F) Volcano plot demonstrating magnitude (log2(fold change)) and significance (-log10(adjusted p-value)) of changes in gene expression from whole-animal RNA sequencing of glial XBP-1s versus wild-type animals at day 2 of adulthood. Differentially expressed genes (DEGs) shown in red (up-regulated, adjusted p-value < 0.05 and log2(fold change) > 0.5, n = 86) and blue (down-regulated, adjusted p-value<0.05 and log2(fold change) < 0.5, n = 22). Labeled genes and DEGs with dark red (up-regulated DEGs) and dark blue (down-regulated DEGs) dots correspond to HLH-30 targets which are labeled with their corresponding gene. (G) Comparison of log2(fold change) of the 86 upregulated DEGs in glial XBP-1s animals compared to wild-type animals (top) compared to the log2(fold change) of these genes in glial XBP-1s animals with a loss-of-function *hlh-30* mutation, *hlh-30(tm1978),* compared to glial XBP-1s animals alone (bottom). log2(fold change) is color coded via a heatmap from warm (up-regulated) to cool (down-regulated). Green dots above the heat map represent HLH-30 target genes. Statistics shown in Table S6.

We next asked if the intestinal nuclear localization of HLH-30 in glial XBP-1s animals is activated in a cell non-autonomous manner or if it is simply a byproduct of UPR^ER^ activation in the periphery. It has previously been shown that glial XBP-1s animals transmit a signal via neuropeptide release and dense core vesicles (DCVs) that is detected by the intestine to activate the UPR^ER^ (Frakes *et al*., 2020). We observed a reversal of the HLH-30::GFP intestinal nuclear localization back to the cytoplasmic localization found in wild type animals when visualizing HLH-30::GFP in glial XBP-1s animals with a loss-of-function mutation that disrupts DCV exocytosis, *unc-31(e928)* (Figure 3D). Importantly, the *unc-31(e928)* mutation did not alter HLH-30 localization, nor did it block the ability of HLH-30 to transit to the nucleus when animals were treated with heat stress, a known activator of HLH-30 (Figure S3A) (Lin et al., 2018).

Glial and neuronal signaling of XBP-1s activation are distinct from each other. Glial XBP-1s animals do not signal to the periphery to activate the UPR^ER^ using neurotransmitters or small clear vesicles (SCV), as in the neuronal XBP-1s paradigm (Frakes *et al*., 2020; Higuchi-Sanabria *et al*., 2020; Ozbey *et al*., 2020; Taylor and Dillin, 2013). To test if peripheral HLH-30 activation is specific to DCV signaling in glial XBP-1s animals, we blocked SCV release via an *unc-13* loss-of-function mutation, *unc-13(e51)*, and found that the loss of SCV release did not suppress the activation of HLH-30 in the periphery (Figure S3B). Taken together, glial XBP-1s animals transmit a distinct signal through DCV release which is detected by the intestine and leads to increased nuclear localization of HLH-30

As a final test if HLH-30 localization was a byproduct of ER stress activation in glial XBP-1s animals, HLH-30::GFP animals were treated with the drug tunicamycin, which induces ER stress via inhibiting protein glycosylation, thus blocking protein folding and transit through the ER. Tunicamycin-induced ER stress does not affect the nuclear localization of HLH-30 (Figure 3E). Furthermore, animals treated with tunicamycin do not activate the transcription of HLH-30 autophagy and lysosomal related target genes (Figure S3C) (Lapierre et al., 2013). Interestingly, animals lacking functional XBP-1, via an *xbp-1(tm2492)* deletion had suppressed nuclear localization of HLH-30 (Figure S3D). Taken together, glial XBP-1s animals transmit a distinct signal to activate HLH-30 in their periphery that cannot be accomplished by cell autonomous ER stress, the signal must come in part from glial cells.

Surprised to find that HLH-30 could be activated cell non-autonomously from glial cells by XBP-1s, we asked whether HLH-30 could activate itself cell non-autonomously if activated in these same glial cells. Ectopic overexpression of *hlh-30* across the entire animal has been shown to increase lifespan and induce HLH-30 target genes (Lapierre *et al*., 2013). Using the same *hlh-30* construct, but now restricted to expression in the four cephalic sheath glia, we did not observe nuclear localization of HLH-30 in peripheral cells (Figure S3E), nor do we find increased lifespan of these animals (Figure S3F). These data indicate that the expression *of xbp-1s* in CEPsh glia can uniquely activate HLH-30 in the periphery and extend lifespan, which cannot be simply achieved by *hlh-30* expression in these cells.

### HLH-30 transcriptionally regulates a high proportion of upregulated genes in glial XBP-1s animals

Finding that HLH-30 is oriented in a unique position to specifically regulate the downstream effects of glial XBP-1s induced longevity, we sought to better understand how HLH-30 could execute these downstream functions in such a distinctive manner. To do so, we asked if the transcriptional and phenotypic changes we observed in the peripheral cells were mediated, in part or completely, by HLH-30.

RNA sequencing revealed substantial gene expression changes in glial XBP-1s animals when compared to wild-type animals, with over a third of upregulated genes being HLH-30 transcriptional targets. To identify HLH-30 target genes that are significantly changed in the glial XBP-1s animals, we generated a list of genes that were previously reported to be HLH-30 targets and had HLH-30 binding sites in the immediate upstream region from the start codon (Grove et al., 2009; Oki et al., 2018; Visvikis et al., 2014). Of the 86 upregulated genes (p < 0.05 and log2(fold change) > 0.5) in glial XBP-1s animals, 36 are HLH-30 targets (Figure 3F, Table S1). These genes have roles in lysosomal lipid and protein catabolism, antioxidant production, immune response, and the UPR^ER^ (Table S2). Additionally, we found that that all glial XBP-1s-upregulated genes were at least partly suppressed by a loss-of-function mutation in *hlh-30*, *hlh-30(tm1978)* (Figure 3G and S3G). These data indicate that constitutive activation of *xbp-1s* in the four cephalic sheath glia in *C. elegans* leads to activation of a peripheral metabolic transcriptional program partly through the activation of HLH-30.

### Glial XBP-1s promotes peripheral autophagy activation

HLH-30 serves critical functions in autophagy through its regulation of numerous autophagy and lysosomal related genes. Because we observed activated HLH-30 in the periphery of glial XBP-1s animals, we hypothesized that there would be an increase in autophagy activation in their distal intestinal cells. First, we asked if glial XBP-1s animals had changes in autophagosome (AP) and autolysosome (AL) content. The number of APs and ALs was determined using the dual-fluorescent reporter of LGG-1, the *C. elegans* ortholog of Atg8, mCherry::GFP::LGG-1 (Chang et al., 2017). LGG-1 is cleaved during AP formation, conjugated to phosphatidylethanolamine, and inserted into the AP membrane. The tandem mCherry/GFP reporter allows for determination of APs versus ALs, due to GFP fluorescence quenching in the highly acidic lysosomal environment (Chang *et al*., 2017). Thus, yellow puncta (co-labeled with GFP/mCherry (pseudo-labeled in magenta) monitor APs, and red puncta monitor the level of ALs within a cell.

Interestingly, the density of APs remains similar at both day 2 and day 5 of adulthood in wild-type and glial XBP-1s animals (Figure 4A and 4B). We see an increase in the density of ALs in glial XBP-1s relative to wild-type animals at both day 2 and day 5 of adulthood (Figure 4A and 4C). Interestingly, we find that the number of APs and ALs in glial XBP-1s animals is dependent on *hlh-30* (Figure S4A and S4B). To corroborate the *in vivo* analysis of autophagy induction, we asked if glial XBP-1s animals had an increase in autophagy and lysosomal genes that are known targets of HLH-30 (Lapierre *et al*., 2013). Indeed, we found a significant increase in the transcripts of *hlh-30*, autophagy-related, and lysosomal-related genes via qPCR (Figure S4C). These data indicate that expression of XBP-1s in CEPsh glia cell non-autonomously activates autophagy peripherally in their intestines over age.

**Figure 4.**
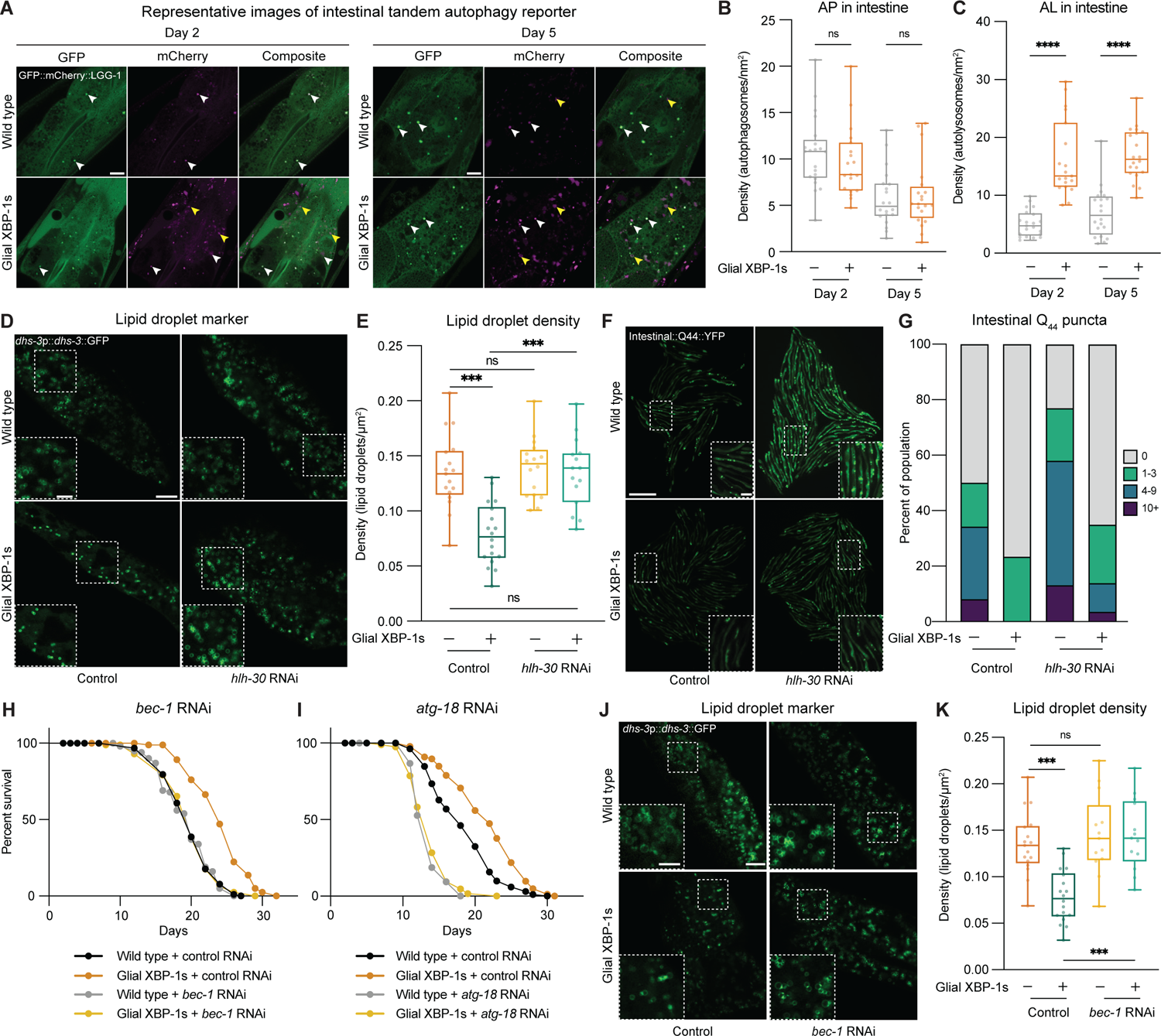
Glial XBP-1s cell non-autonomously activates peripheral autophagy, which is required for lifespan extension and reduced lipid levels. (A) Representative Airyscan micrographs from intestine of day 2 and day 5 wild-type and glial XBP-1s transgenic animals expressing the tandem autophagy reporter *mCherry::GFP::lgg-1* (green, GFP, magenta, mCherry). White arrowheads indicate autophagosomes, yellow arrowheads indicate autolysosomes. Scale bar, 10 µm. (B-C) Quantification of mCherry puncta co-localized with GFP (autophagosomes (AP)) (B) or containing mCherry alone (autolysosome (AL)) (C) in the intestine of wild-type and glial XBP-1s transgenic animals at day 2 and day 5 of adulthood. Data are from three independent experiments, each with ≥ 5 animals. Statistics by Kruskal-Wallis with Dunn’s multiple comparison test, p < 0.0001 (****). (D) Representative Airyscan micrographs of *dhs-3*p::*dhs-3*::GFP labeled lipid droplets from the intestine of day 2 wild-type and glial XBP-1s adult animals grown on HT115 *E. coli* expressing control or *hlh-30* RNAi. Scale bar, 10 µm. Scale bar of inset, 5 µm. (E) Changes in lipid droplet density of wild-type and glial XBP-1s animals grown on control or *hlh-30* RNAi from (D). Box plot shows median, whiskers are minimum to maximum values, each dot is representative of one animal. Statistics by Kruskal-Wallis with Dunn’s multiple comparison test, p < 0.001 (***), p > 0.05 (ns). N > 15, and from three independent replicates. (F) Representative fluorescent micrographs of age-dependent accumulation of transgenic polyQ_44_-YFP aggregates in wild-type or glial XBP-1s animals grown on control or *hlh-30* RNAi. Scale bar, 1 mm. Scale bar of inset, 200 µm. (G) Quantification of age-dependent polyQ_44_ aggregates from (F), grouped into animals with 0 (gray), 1-3 (green), 4-9 (blue), or >10 (purple) aggregates. N > 100 per condition. Data is representative from three independent replicates. Statistics done by Chi-squared test for independence with adjusted residual and Bonferroni correction, statistics shown in table S5. (H-I) Survival of wild-type and glial XBP-1s animals on control, *bec-1* (D) or *atg-18* RNAi (E) at 20°C. Lifespan is representative of three independent replicates. See Table S1 for lifespan statistics. (J) Representative Airyscan micrographs of transgenic *dhs-3*p::*dhs-3*::GFP labeled lipid droplets from the intestine of day 2 wild-type and glial XBP-1s adults grown on control or *bec-1* RNAi. Scale bar, 10 µm. Scale bar of inset, 5 µm. (K) Changes in lipid droplet density of wild-type and glial XBP-1s animals grown on control or *bec-1* RNAi from (J). Box plot shows median, whiskers are minimum to maximum values. Statistics by Kruskal-Wallis with Dunn’s multiple comparison test, p < 0.001 (***). N > 15 per condition from three independent replicates.

While it is of great interest to determine if the activation of peripheral autophagy is dependent on DCV release, using the *unc-31(e928)* mutant to probe this provides some challenges. First, mutations in *unc-31* are known to disrupt neuropeptide signaling, including insulin-like signaling, which in turn can activate autophagy in these animals (Ailion et al., 1999; Hansen et al., 2008). We confirmed this probing for targets of HLH-30 in *unc-31(e928)* mutant animals and found that there was significant upregulation of these genes in *unc-31(e928)* mutant animals (Figure S4E).

### HLH-30/TFEB is required for the reduction in lipid droplets and intestinal protein aggregates in glial XBP-1s animals

Since HLH-30 has transcriptional control over genes involved in lysosome biogenesis and lipid catabolism, we asked if the decreased lipid droplet content in the intestine of glial XBP-1s animals is dependent on *hlh-30*. We found that *hlh-30* is required for reduction of lipid droplets in glial XBP-1s animals (Figure 4D). When *hlh-30* is knocked down in glial XBP-1s animals the density of intestinal lipid droplets returns to wild-type levels (Figure 4E). These data point to a requirement for *hlh-30* in the reduction of lipid stores in the intestines of glial XBP-1s animals.

HLH-30 has also been shown to be necessary for protection against proteotoxicity in *C. elegans* overexpressing *xbp-1s* pan-neuronally (Imanikia *et al*., 2019a). It has also been shown that expression of *xbp-1s* in CEPsh glia leads to a significant reduction in aggregation of yellow fluorescent protein (YFP)-tagged polyglutamine (polyQ) repeat expansion protein in the intestine (Frakes *et al*., 2020). Proteins with polyQ expansions are implicated in neurodegenerative disorders, such as Huntington’s disease, and when expressed in *C. elegans* can model such protein-folding disorders and serve as a proxy for proteostasis capacity (Lieberman et al., 2019; Morley et al., 2002). We examined the number of intestinal polyQ-YFP aggregates in adulthood and found that the reduction of aggregates in glial XBP-1s animals is dependent on *hlh-30* (Figure 4F and 4G, S4D). These data indicate that the glial XBP-1s paradigm protects against intestinal polyQ aggregation through the activity of HLH-30.

### Macroautophagy is required for lifespan extension and lipid droplet depletion in glial XBP-1s animals

The ability to activate autophagy declines with age and is increased in numerous conserved longevity paradigms (Chang *et al*., 2017). We observed an increase in the expression of autophagy genes (Figure S4C) and increased autophagic capacity in glial XBP-1s animals (Figure 4A-4C). Therefore, we tested if autophagy genes are required for the lifespan extension of glial XBP-1s animals. Knockdown by RNAi of autophagy genes that are required for the induction of macroautophagy, *bec-1* (autophagosome membrane nucleation) and *atg-18* (phosphoinositide 3-phosphate binding), abrogated the lifespan extension of glial XBP-1s animals (Figure 4H and 4I). These data indicate that macroautophagy is required for lifespan extension in glial XBP-1s animals.

Lipid catabolism is driven through lipophagy, the autophagic degradation of lipid droplets, and lipolysis in lysosomes. We found an expansion of lysosomes, upregulation of HLH-30 lipid catabolism genes, and a dependence on *hlh-30* of the lipid depletion in the intestines of glial XBP-1s animals. Therefore, we hypothesized that this depletion is caused by macroautophagy of lipid droplets. We find that knockdown of *bec-1* fully suppresses the lipid droplet depletion found in glial XBP-1s animals (Figure 4J and 4K).

### Altered ER structures in glial XBP-1s animals are dependent on macroautophagy genes and co-localize with autophagosomes and autolysosomes

After seeing changes in autophagic activation concurrent with HLH-30 activation, we asked whether the changes in ER morphology detected in glial XBP-1s animals (Figure 1I and 1J) are dependent on autophagy (Bernales *et al*., 2006). ERphagy, the selective degradation of the endoplasmic reticulum by autophagy, plays a major role in maintaining ER homeostasis and is involved in the recovery from ER stress (Hubner and Dikic, 2020). We found that the formation of the ER structures seen in glial XBP-1s animals is dependent on macroautophagy in the intestine. Knockdown of core components of the macroautophagy machinery, *atg-18*, *vps-34* (kinase that recruits machinery to form autophagosomes)*, bec-1*, and *lgg-1* (involved in phagophore elongation and cargo recruitment) resolved the formation of the ER puncta detected in glial XBP-1s animals (Figure 5A and 5B). Intriguingly, when determining the dependence on the ER puncta formation on neuropeptide, via *egl-3(ok979)* mutation or DCV release, via *unc-31(e928)* mutation, we found that knock-out of these genes did not suppress the formation of the ER puncta in glial XBP-1s animals (Figure S4F).

**Figure 5.**
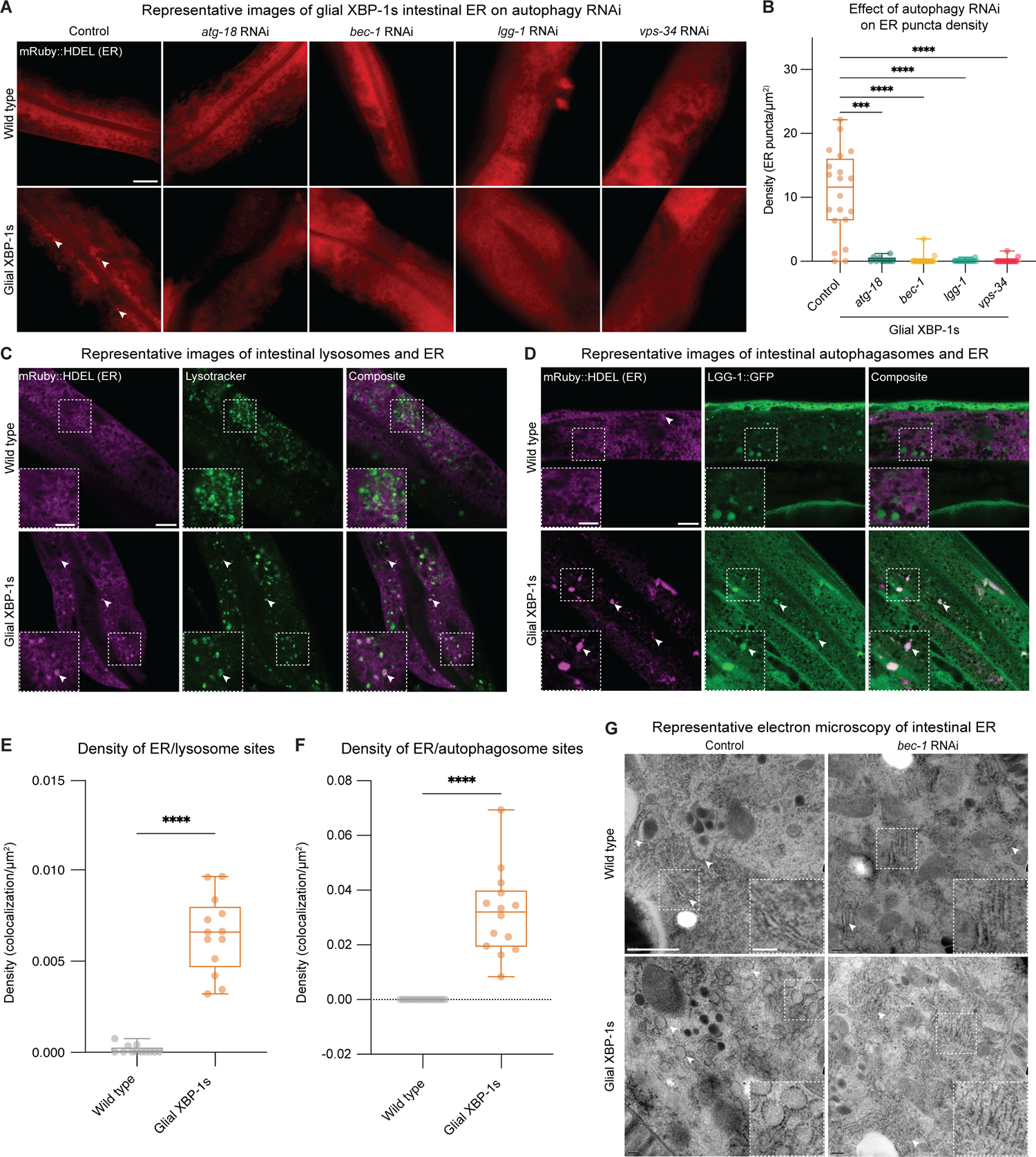
Macroautophagy is required for changes in intestinal ER morphology and lipid depletion in glial XBP-1s animals. (A) Representative Airyscan micrographs from intestine of day 2 wild-type and glial XBP-1s animals transgenic for the ER marker *vha-6p*::ERss::mRuby::HDEL, animals were grown on control or the corresponding RNAi. ER puncta is denoted by arrowheads. (B) Quantification of ER puncta from (A). Density was determined by counting the number of ER puncta per area of intestine expressing *vha-6p*::ERss::mRuby::HDEL. N > 15 per condition from three independent replicates. Statistics by Kruskal-Wallis with Dunn’s multiple comparison test, p < 0.001 (***), p < 0.0001 (****). (C) Representative Airyscan micrographs from intestines of day 2 wild-type and glial XBP-1s animals transgenic for the ER marker *vha-6p*::ERss::mRuby::HDEL (pseudo-colored in magenta) and stained with Lysotracker Blue DND-99 (pseudo-colored in green). Arrowheads mark colocalization. Scale bar, 10 µm. Scale bar of inset, 5 µm. (D) Representative Airyscan micrographs from intestine of day 2 wild-type and glial XBP-1s animals transgenic for the ER marker *vha-6p*::ERss::mRuby::HDEL (magenta) and autophagosome marker LGG-1::GFP (green). Arrowheads mark colocalization. Scale bar, 10 µm. Scale bar of inset, 5 µm. (E) Quantification of density of ER puncta and lysotracker colocalization. Statistics by Mann-Whitney test, p < 0.0001 (****). N > 15 per condition from three independent replicates. (F) Quantification of density of ER puncta and autophagosome colocalization. Statistics by Mann-Whitney test, p < 0.0001 (****). N > 15 per condition from three independent replicates. (G) Electron micrographs of intestine from wild-type and glial XBP-1s animals at day 2 of adulthood grown on either empty vector control or *bec-1* RNAi. Imaging was replicated in triplicate for a total of 15-20 animals being imaged per condition. Arrowheads mark rough endoplasmic reticulum. Scale bar, 1µm. Scale bar of inset, 0.2 µm.

We next investigated if these ER puncta colocalized with markers for lysosomes and autophagosomes. Staining for lysosomes using LysoTracker, a pH-dependent dye which accumulates in acidic lysosomes, in the mRuby::HDEL (ER) strains showed significant colocalization of lysosomes with mRuby::HDEL puncta in glial XBP-1s compared to wild-type animals (Figure 5C and 5E). Using a strain expressing GFP-tagged LGG-1, a protein that localizes to autophagosomal membranes, we found that glial XBP-1s animals have significant colocalization of mRuby::HDEL (ER) and LGG-1::GFP positive puncta (Figure 5D and 5F). We next asked if the circular ER structures we visualized using high-magnification transmission electron microscopy (Figure 1I) were dependent on autophagy activation. Strikingly, we found that knockdown of *bec-1* in glial XBP-1s resolved the formation of these structures (Figure 5G and Figure S4G). Taken together, these results suggest that *xbp-1s* expression in CEPsh glia leads to cell non-autonomous restructured distal ER, which is dependent on macroautophagy components and colocalizes with autophagosomes and lysosomes, indicative of ERphagy activation.

## DISCUSSION

In this work, we have found that CEPsh glial cells of *C. elegans* coordinate peripheral metabolism, autophagy activation, and ER remodeling when the UPR^ER^ is activated in these four cells. Ectopic expression of constitutively active *xbp-1s* in CEPsh glia leads to significant lipid depletion and ER remodeling in the intestine. We find that the transcription factor HLH-30 has significant nuclear localization in intestinal tissue and partly controls expression of a third of genes upregulated in glial XBP-1s animals (Figure 6). Interestingly, mutations in genes required for DCV exocytosis (*unc-31*), suppressed this cell non-autonomous activation of HLH-30 in glial XBP-1s animals. *hlh-30* is required for the significant lifespan extension, reduction in lipid content, and increased intestinal proteostasis found in glial XBP-1s animals. We also find that ectopic *xbp-1s* expression leads to cell-non autonomous activation of autophagy in the periphery, and that autophagy regulators are required for lifespan extension and lipid depletion. Lastly, the ER morphological changes seen in glial XBP-1s animals are dependent on macroautophagy, colocalize with autophagosomes and lysosomes, and may be suggestive of ER-phagy occurring peripherally. Taken together, these data describe a novel role of communication from UPR^ER^ activation in glial cells to the periphery which regulates organismal metabolism and autophagy activation that cannot be achieved by cell autonomous UPR^ER^ activation.

**Figure 6.**
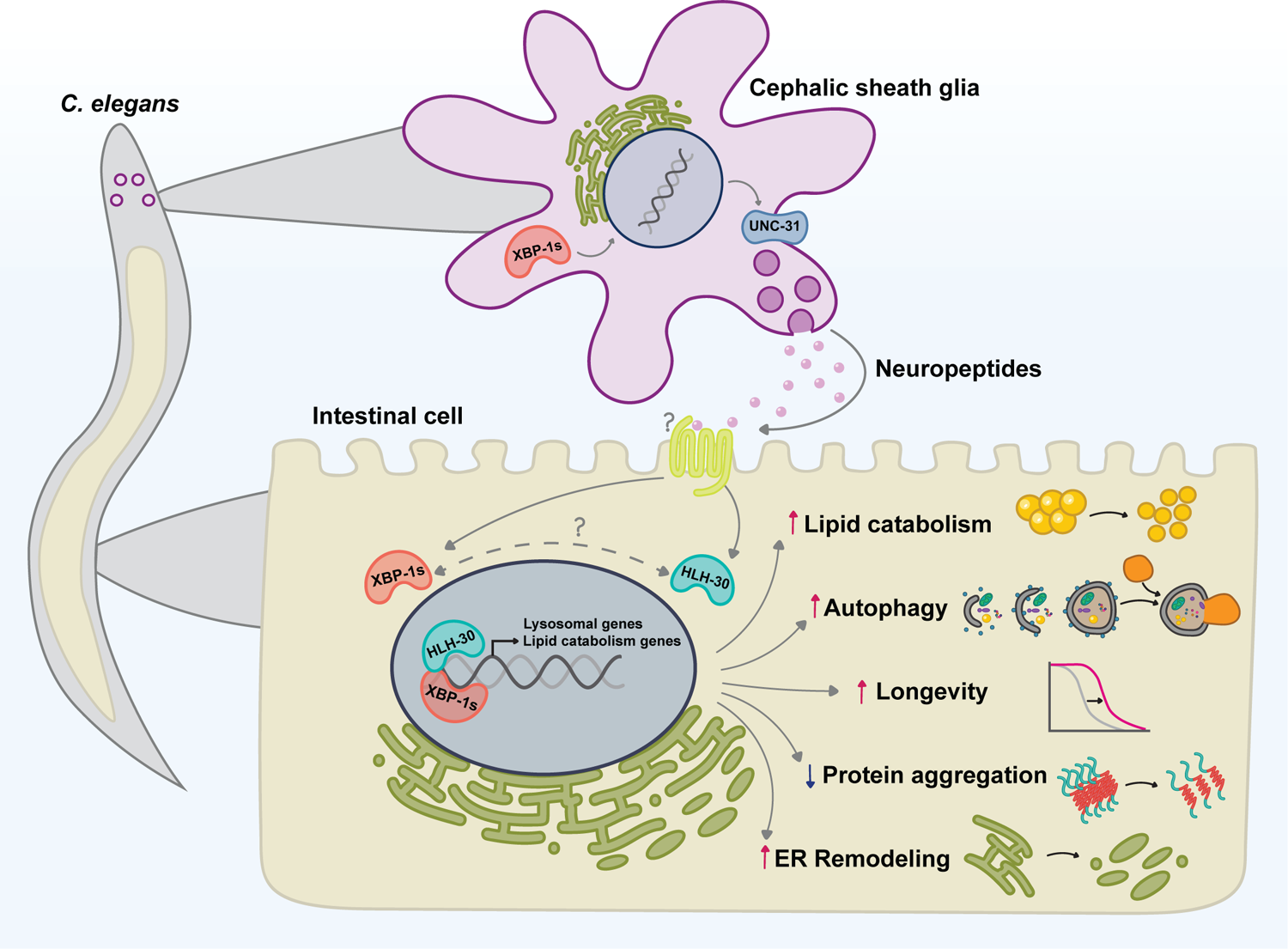
Schematic of glial XBP-1s effects on peripheral metabolism and autophagy. Expression of *xbp-1s* in the four CEPsh glia induces activation of the UPR^ER^ and the transcription factor HLH-30/TFEB in the periphery, reliant on exocytosis of dense core vesicles (*unc-31*). HLH-30 regulates the transcription of lipid catabolism and lysosomal biogenesis genes in glial XBP-1s animals. HLH-30 is required for increased lipid catabolism, activated autophagy, decreased intestinal protein aggregation, lifespan extension, and ER remodeling in glial XBP-1s animals. Autophagy induction is also required for the longevity, lipid catabolism, and ER remodeling phenotypes in the glial XBP-1s paradigm. These data suggest that glia provide cell non-autonomous control over peripheral autophagy and metabolic adaptation during stress.

This work has shown that glial cells have crucial roles in the regulation of organismal autophagy and lipid metabolism beyond the nervous system. Previous work has shown that pan-neuronal ectopic expression of XBP-1s in *C. elegans* regulates cell non-autonomous UPR activation in the gut to extend lifespan and increase stress resistance (Taylor and Dillin, 2013). Communication between neurons and the intestine depends upon SCV release of neurotransmitters, improving lysosome activity and lipid metabolism (Daniele *et al*., 2020; Higuchi-Sanabria *et al*., 2020; Imanikia *et al*., 2019a; Imanikia *et al*., 2019b; Ozbey *et al*., 2020). In contrast, glial cell non-autonomous activation of the UPR^ER^ and lifespan extension is dependent on neuropeptides released from dense core vesicles (DCV) (Frakes *et al*., 2020). Additionally, we find DCV-dependent activation of HLH-30 in glial XBP-1s animals, in contrast to neuronal XBP-1s animals that do not activate HLH-30 peripherally (Imanikia *et al*., 2019a). These data suggest that glial cells communicate to the periphery to coordinate the induction of lipid catabolism and lysosome activation through a mechanism independent and distinct from that of neurons, which has been found to be dependent on the neurotransmitters tyramine, acetylcholine, serotonin, and dopamine (Higuchi-Sanabria *et al*., 2020; Ozbey *et al*., 2020). Further studies into the origin and nature of this glial signal, which in *C. elegans* may act as a primitive hormone signal, are of great interest. Discovery of the relevant glial signal would identify therapeutic interventions to potentially reverse age-onset proteome and autophagic decline seen in neurodegenerative and metabolic diseases.

The transcription factor TFEB/HLH-30 controls lysosomal biogenesis and autophagy by regulating the expression of genes in the coordinated lysosomal expression and regulation (CLEAR) network (Settembre *et al*., 2011). Interestingly, knockout of *Tfeb* in hepatocytes of mice leads to impaired lipid degradation pathways in these cells (Settembre *et al*., 2013). Conversely, overexpression of *Tfeb* in liver cells of mice fed a high-fat diet or in obese mice, caused by a homozygous mutation in the gene responsible for the production of leptin (*ob/ob*), a hormone responsible for regulating energy balance by inhibiting hunger, attenuates the development of obesity (Settembre *et al*., 2013). In *C. elegans*, HLH-30 couples nutrient availability to transcription of lysosomal lipases and autophagy, controlling fat storage and aging (O’Rourke and Ruvkun, 2013). It was recently found that XBP1s regulates autophagy in murine liver cells through the activation of TFEB. Overexpression of *Xbp-1s* enhanced TFEB transcription and autophagy, while overexpression of *Tfeb* ameliorated the glucose intolerance seen in *Xbp1* liver knockout mice with diet-induced obesity (Zhang et al., 2021). Our findings implicate numerous genes being co-regulated by both XBP-1s and HLH-30 in these glial XBP-1s animals. Further investigation into the XBP-1s/HLH-30 axis and how these pathways converge in metabolic health and dysfunction is warranted.

The failure of the UPR^ER^ to function properly is a hallmark of numerous age-related neurodegenerative and metabolic disorders. In states of chronic overnutrition, the UPR^ER^ machinery can be overwhelmed, leading to persistent and unresolved ER stress. Mice that are heterozygous for a null XBP-1 allele develop glucose intolerance, severe insulin resistance, and diabetes (Ozcan et al., 2004). Additionally, *ob/ob* mice have impaired UPR^ER^ signaling and decreased nuclear localization of XBP-1s in their livers (Park et al., 2014). Interestingly, mice with a conditional deletion of *Xbp-1* in neurons and glia are more susceptible to diet-induced obesity and become leptin resistant (Ozcan et al., 2009). Surprisingly, activation of XBP-1s in hypothalamic pro-opiomelanocortin (POMC) neurons of mice leads to cell non-autonomous induction of the UPR^ER^ in the liver which protects the mice from diet-induced obesity and helps maintains metabolic homeostasis (Williams *et al*., 2014). Cell non-autonomous activation of the UPR^ER^ from neurons to the periphery has also been seen in physiological settings, food perception alone in mice induces global metabolic changes involving systemic UPR^ER^ activation (Brandt *et al*., 2018). This cell non-autonomous signaling of the UPR^ER^ points to a conserved mechanism of global control of organismal metabolic homeostasis by neural UPR^ER^ activation. Interrogating the potential role of mammalian glial XBP-1s as a therapeutic target in these chronic metabolic disorders is therefore of great interest.

Because glia have critical roles in providing metabolic support for neurons, we speculate that they are well-positioned to integrate the metabolic state of the brain and mediate communication with peripheral tissues. It is hypothesized that glial cells first evolved as static support cells for neurons, but with increasing nervous system complexity their roles shifted to increase neural function capacity and to generate efficient energy metabolism in the nervous system (Hartline, 2011; Rey et al., 2021) There are numerous subpopulations of glial cell types that are positioned at major cruxes of metabolism, including astrocytes in the hypothalamus, tanycytes in contact with both the cerebral spinal fluid and the arcuate nucleus of the hypothalamus, and enteric glia in the gut-brain axis (García-Cáceres et al., 2016; Porniece Kumar et al., 2021; Seguella and Gulbransen, 2021). In humans, the brain is the primary consumer of glucose, using a fifth of total ATP stores, primarily from glucose in blood circulation, with glial cells playing a critical role in the transport of glucose (Erbsloh et al., 1958). In the CNS, glial cells uptake glucose and generate glycolysis-derived lactate which is transported into neurons where it is then metabolized to acetyl coenzyme A, a key energy input into the tricarboxylic acid cycle (Liu et al., 2017). Additionally, two separate studies have shown that insulin signaling in hypothalamic astrocytes and tanycytes is required for brain glucose sensing and systemic glucose metabolism (Garcia-Caceres et al., 2016; Porniece Kumar *et al*., 2021). Further identification of the processes that are specific to glial XBP-1s regulation of neuronal and systemic metabolic homeostasis is warranted.

Here, we show that glia play a critical role in coordinating peripheral metabolic homeostasis and autophagy activation via XBP-1s activation and cell non-autonomous activation of HLH-30. Activation of these processes relies solely on activation of *xbp-1s* in CEPsh glia. Neither HLH-30 overexpression in glial cells or cell autonomous induction of the UPR^ER^ can turn on these processes in the periphery. We believe that glial communication of ER stress is an advantageous organismal adaptation by which detection of ER stress can be sensed and systemically transmitted to promote longevity, stress resistance, and metabolic health. Our work furthers the argument of the UPR^ER^ being a major sensor and coordinator of organismal homeostasis and highlights the important roles of glial cells in coordinating systemic metabolism and proteostasis. The evolution of this process in metazoa ensures communication of stress and metabolism across cell and organ systems to promote health of the entire organism.

## ACKNOWLEDGEMENTS

We thank the Caenorhabditis Genetics Center (CGC) (P40 OD010440) for several strains used in this study and the Dillin lab for comments and discussion throughout the project. M.G.M is supported by NIH (F31 AG060660). A.E.F. was supported by the NIH (F32 AG051355). A.D. is supported by the Thomas and Stacey Siebel Foundation, the Howard Hughes Medical Institute, and NIH grants R37 AG024365, R01 ES021667, R01 AG055891 and R01 AG059566.

## AUTHOR CONTRIBUTIONS

M.G.M conceived the project. M.G.M, S.M, A.S., J.D., A.E.F., M.V., C.M., A.F., and M.S. performed experiments. M.G.M, S.M, A.E.F., A.S., M.V., and A.F. analyzed data. A.D. supervised the project. M.G.M. wrote the manuscript with input from all authors.

## DATA AND MATERIALS AVAILABILITY

The raw RNA-seq data are on the NCBI short read archive. Further information and requests for resources and reagents should be directed to and will be fulfilled by the lead contact, Dr. Andrew Dillin.

## METHODS

### *C. elegans* strains and details

Nematodes were maintained at 15°C or 20°C on standard nematode growth medium (NGM) agar plates seeded with *Escherichia coli* (*E. coli*) OP50. All experimentation was performed at 20°C on OP50 or HT115 *E. coli* K12 strain containing pL4440 empty vector control or expressing double stranded RNA containing the sequence of the target gene for RNA interference (RNAi) experiments. RNAi strains were all isolated from Vidal or Ahringer libraries and sequence verified before use. For all experiments, animals are synchronized using a standard bleaching protocol where carcasses of reproductive animals are degraded using a bleach solution (1.8% sodium hypochlorite, 0.375 M KOH), followed by 5 washes with M9 buffer (22 mM KH_2_PO_4_ monobasic, 42.3 mM NA_2_HPO_4_, 85.6 mM NaCl, 1 mM MgSO_4_) and eggs were then platted onto NGM plates spotted with OP50. Glial XBP-1s animals have a slight developmental delay, to account for this in glial XBP-1s were bleached 8 hours before wild type animals for all experiments. All strains used in this study are listed in Table S4.

Transgenic worms (glial HLH-30) were synthesized by injecting N2 worms with plasmid pMM26 listed in Key Resources Table at 50ng/ul with co-injection marker pEK2 (unc-122p::RFP (coelomocyte RFP)) at 50ng/ul and 100ng/ul of pD64 vehicle as filler DNA. *hlh-30* genomic DNA includes isoforms A,F,G, and H. Worms positive for *unc-122*p::RFP were selected for stable arrays.

### Lifespan analysis

Lifespan analyses were performed on NGM plates at 20°C on HT115 *E. coli* strain expressing empty vector control or the designated double-stranded RNAi. Animals were age-synchronized by bleaching and plating on HT115 and grown from hatch to adulthood at 20°C, unless otherwise stated. Bleach times were determined based on developmental delay of strains, so that all animals reach day 1 of adulthood at the same time. Adult nematodes were manually moved away from progeny onto fresh plates every day or every other day during reproduction. Nematodes were scored every 1-2 days for death or censored due to bagging, vulval explosions, or crawling off the plate. Lifespan analyses were blinded and performed a minimum of 3 times by at least 2 independent researchers. Representative data from one of the 3 trials were presented as a Kaplan-Meier curve. Prism 9 was used for statistical analyses, p-values were calculated using long-rank (Mantel-Cox) method.

### Stereomicroscopy for fluorescent reporters

Animals were age-synchronized by bleaching, eggs were plated on NGM plates spotted with OP50 *E. coli* or HT115 *E. coli* expressing empty vector control or the designated double-stranded RNAi and grown to adulthood at 20°C. Animals were moved onto unspotted NGM plates at day 1 or day 2 of adulthood and anesthetized with 5 µL 100 mM sodium azide. Animals were imaged using a Leica M250FA automated fluorescent stereoscope equipped with a Hamamatsu ORCA-ER camera, standard GFP filter, and driven by LAS-X software. Each microscopy experiment was performed a minimum of 3 independent times.

### BODIPY 493/503 staining

Animals were age-synchronized by bleaching, eggs were plated on NGM plates spotted with OP50 bacteria. At day 2 of adulthood, animals were washed off using M9, spun down at 1,000 RCF, and washed 3X with M9 in 15mL conical tubes. M9 was aspirated and replaced with 500ul of 4% paraformaldehyde (Electron Microscopy Sciences). Animals were fixed for 15 minutes while rotating in the dark at 20°C. Animals were then frozen in liquid nitrogen and thawed in a 37°C water bath, followed by wash with 1X PBS, spin down at 1,000 RCF and repeated a total of 3 times. 10 ul of 1mg/mL BODIPY 493/503 (Invitrogen) was diluted 1:1000 in 10mL of PBS. Animals then were put on a rotator at 20°C in the dark for one hour. After incubation with BODIPY 493/503, animals were washed 3 times with 1X PBS and spun down at 1,000 RCF after each wash. 5 mL 1X PBS was placed into each tube and fixed animals were incubated overnight at 4°C in the dark. The next day animals were washed 3 times with 1X PBS and then run through the COPAS BioSorter for whole worm fluorescence quantification.

### Imaging of HLH-30 nuclear translocation assay

Day 2 adult animals were anesthetized in 15 µL 100 mM sodium azide on microscope slides and covered with a cover slip fastened with nail polish. Worms were imaged using a Zeiss AxioObserver.Z1 microscope equipped with a Lumencor SOLA light engine and a ZEISS Axiocam 506 camera, driven by ZEISS ZEN Blue software using a 63x/1.4 Plan Apochromat objective. Images of the anterior 30% of the intestine were taken using a 63X objective/1.4 Plan Apochromat objective and standard GFP filter (ZEISS filter set 46 HE). To avoid potential translocation phenotypes caused by exposure to the mounting and imaging conditions, all scoring was conducted within 5 minutes after mounting.

### Whole animal fluorescence quantification

For large-scale quantification of fluorescent and dyed animals, a Union Biometrica complex object parameter analysis sorter (COPAS) BioSorter was used to measure GFP fluorescence of individual worms for quantification. Animals were washed off NGM plates using M9 wash, and run through the COPAS BioSorter using a 488nm light source. Total integrated GFP fluorescence was normalized to extinction (thickness) of each individual animal. Data was processed by censoring events that reached the maximum peak height for green or extinction values (PH Green and PH Ext = 65532) and censoring events <300 and >1000 time of flight (length) to remove progeny and greater than one worm per event. Values of green <=10 were removed for strains with visible fluorescence (BODIPY 493/503, *dhs-3*p::*dhs-3*::GFP) as these values are most likely air bubbles detected by the sorter. All data was normalized to the mean of fluorescent values of wild-type animals. Each data point represents an individual animal measurement. All experiments were performed a minimum of 3 independent times. Statistics were calculated using Prism 9 software.

### Measurement of polyQ aggregates

Animals were age-synchronized by bleaching and adding eggs to NGM plates spotted with HT115 *E. coli* expressing empty vector control (pL4440) or the designated double-stranded RNAi and grown to adulthood at 20°C. Animals were moved at day 2 of adulthood to new NGM plates spotted with HT115 *E.coli* and away from their progeny. Fluorescent microscopy was performed at day 3 of adulthood. Images were blinded and YFP-positive puncta were counted per animal, for a minimum of 100 animals per condition. Fluorescent microscopy is representative of 3 independent experiments.

### qPCR

Animals were grown on standard NGM plates spotted with OP50 from hatch at 20°C until day 1 of adulthood. Animals were collected by washing off with M9 buffer. M9 was subsequently aspirated and replaced with Trizol (Invitrogen). Animals were homogenized via a freeze-thaw process 3x with liquid nitrogen. Following the final freeze-thaw cycle, chloroform was added at a 1:5 volumetric ratio (chloroform:Trizol) for aqueous separation of RNA via centrifugation in heavy gel phase-lock tubes (VWR). Aqueous phase was transferred into isopropanol followed by RNA purification using a Qiagen RNeasy Mini Kit as per manufacturer’s instructions. 0.5-2 ng of purified RNA was used for cDNA synthesis using the Qiagen QuantiTect Reverse Transcriptase kit as per manufacturer’s instructions. qPCR analysis was performed using a general standard curve protocol using SYBR-green on an Applied Biosystems QuantStudio 384-well qPCR machine. Per sample, 3 biological replicates and 4 technical replicates per independent biological replicate were run.

For the tunicamycin qPCR experiment, animals were synchronized and grown to D1 of adulthood on EV RNAi plates. Animals were then treated with either 1% DMSO or tunicamycin (25 ng/µl), rotating at 20°C for 1 hour in a 15 ml conical tube. After an hour of treatment, M9 was aspirated, and animals were washed 3X with M9 buffer.

### RNA isolation, sequencing, and analysis

Animals were bleach synchronized and grown to day 2 of adulthood on large NGM plates spotted with 1mL of OP50. Around 1,000 animals per condition and replicate were collected by washing off with M9 buffer. Each condition had a total of 5 biological replicates. Animals were spun down at 1,000 RCF for 30 seconds, M9 removed and washed again for a total of 3 washes. M9 was then replaced with 1 mL of Trizol (Invitrogen) and animals were subsequently froze, via liquid nitrogen, and thawed, in a 37°C water bath, 3 times. After the final thaw, chloroform was added at a ratio of 1:5 (chloroform:trizol) for aqueous separation of RNA using heavy gel phase-lock tubes (VWR, 10847-802). Library preparation and RNA-sequencing was performed by Genewiz (Azenta Life Sciences). Library preparation was performed with Poly A selection and HiSeq xSequencing using NEBNext Ultra RNA Library Prep Kit for Illumina following manufacturer’s instructions (NEB, Ipswich, MA, USA). Sequencing libraries were clustered onto 1 lane of a flow cell, loaded on Illumina HiSeq instrument (4000 or equivalent) and samples were sequenced using 2×150bp Paired End configuration. Base calling was conducted on HiSeq Control Software. For RNA-sequencing analysis, raw sequencing data were uploaded to the Galaxy project web platform and the public server at usegalaxy.org was used to analyze the data (Afgan et al., 2016). Paired end reads were aligned using the Kallisto quant tool (version XXX) with WBcel235 as the reference genome. Fold changes and statistics were generated using the DESeq2 tool with Kallisto quant count files as the input. The fold change and adjusted-p values generated by the DESeq2 analysis were used to plot the data using GraphPad Prism 9.

### Electron microscopy imaging and analysis

Whole animal samples were processed for transmission electron microscopy (TEM) as previously described (Klapper *et al*., 2011). Briefly, staged animals were subjected to high-pressure freezing (BAL-TEC HPM 010) and freeze-substituted with acetone/resin series (25%-50%-75%-100% resin). Resin-cured animals were sectioned into 70nm sections and imaged using a FEI Tecnai 12 transmission electron Microscope on formvar-coated mesh grids.

TEM images were analyzed using ImageJ. Rough ER was determined as described (Sanvictores and Davis, 2022); the rough ER is decorated with ribosomes which are detected in the TEM images as black dots. Using the freehand selection tool, the area of rough ER cisternae was determined as the space between the black ribosome dots which outline the rough ER cisternae. The circularity index was calculated using the circularity measurement in ImageJ. The circularity index was determined using the following formula *circularity = 4pi(area/perimeter^2)*.

### Airyscan microscopy

Live animals at their listed respective age were paralyzed in 15 µL sodium azide (100 mM) on microscope slides and covered with a cover slip fastened with nail polish. Fluorescent images were obtained with a Zeiss LSM900 Airyscan microscope. Images were quantified using ImageJ (FIJI).

### LysoTracker Blue DND-22 staining

Animals were grown from hatch to day 2 of adulthood on OP50 *E. col* seeded NGM plates and then moved to OP50 seeded plates containing 10 µM ThermoFisher scientific LysoTracker Blue DND-22 (Invitrogen) (1mM stock diluted 1:100) or an equivalent volume of M9 buffer for 2 hours. After 2 hours worms were moved and placed on OP50 plates for 1 hour. Worms were then imaged by AiryScan microscopy, using 10 animals per genotype and 3 independent biological replicates. Quantification was carried out using ImageJ.

### Pharyngal pumping assay

Animals were grown to day 1 on NGM plates seeded with OP50 *E. coli*. Young adults (early day 1) were then transferred to a new NGM plate seeded with OP50 *E. coli* and allowed to crawl on the plate for 15 min. Pharyngeal pumping rate was determined by visual counting of the movement of the grinder under a dissecting microscope for a total of 60 seconds, counting for 10 seconds with a 10 second break in between.

### Heat shock assay

Animals were grown to day 2 of adulthood on NGM plates spotted with OP50 *E. coli* and exposed to a heat shock for 3 hours at 35°C. HLH-30::GFP nuclear localization in the intestine was scored immediately as described in the ‘Imaging of HLH-30 nuclear translocation assay’ methods section.

### Tunicamycin assay

Animals were grown to day 1 of adulthood, washed with M9 buffer for a total of three times, and then treated with either tunicamycin (25 ng/µl) or DMSO (1%) diluted with M9 buffer in a 15 mL conical tube. The animals were left rotating at 20°C in the dark for 1 hour. The animals were then washed 3 times with M9 buffer and plated on OP50 *E. coli* spotted NGM plates to recover and imaged 1 hour afterwards.

## SUPPLEMENTAL FIGURE LEGENDS

**Figure S1.**
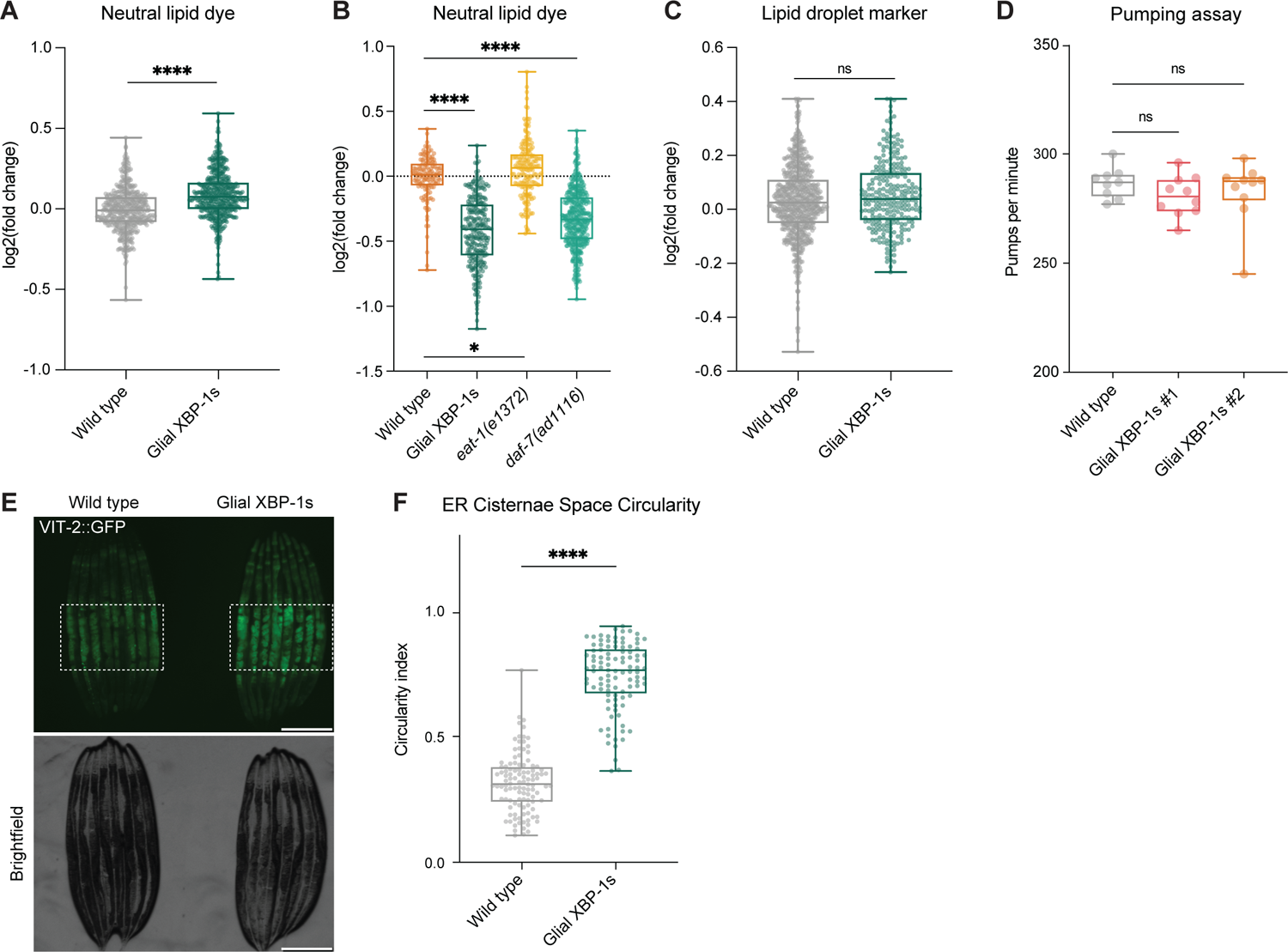
(A) Quantification of fixed, unstained wild-type and glial XBP-1s animals using a COPAS BioSorter to determine whole worm autofluorescence quantification. Animals were grown to day 2 of adulthood on OP50 bacteria. Intensity of green values was compared to mean of fixed, unstained wild-type animals to determine fold change. Box plot shows median, whiskers are minimum to maximum values. N > 100 animals per condition. Statistics by Mann-Whitney test, p < 0.0001 (****). (B) Quantification of wild-type, glial XBP-1s, *eat-2*(*e1372*), and *daf-7*(*ad1116*) animals at day 2 of adulthood stained with BODIPY 493/503 using a COPAS BioSorter for whole worm BODIPY 493/503 dye quantification. BODIPY 493/503 staining intensity was compared to mean of wild-type animals to determine fold change. Box plot shows median, whiskers are minimum to maximum values. N > 100 animals per condition. Statistics by One-way ANOVA with Sidák’s multiple comparison test, p < 0.0001 (****), p < 0.5 (*). (C) Quantification of wild-type and glial XBP-1s using a COPAS BioSorter to determine whole worm autofluorescence quantification. Animals were grown to day 2 of adulthood on OP50 bacteria. Intensity of autofluorescence was compared to mean of wild-type animals to determine fold change. Box plot shows median, whiskers are minimum to maximum values. N > 100 animals per condition. Statistics by Mann-Whitney test, p > 0.05 (ns = not significant). (D) Pharyngeal pumping per minute of wild-type, glial XBP-1s (integrated strain #1), and glial XBP-1s (integrated strain #2) animals at the young adult stage. The graph shows representative data of two experiments, n = 10 animals per condition. Statistics by Kruskal-Wallis with Dunn’s multiple comparison test, p > 0.05 (ns). (E) Representative fluorescent micrograph of wild-type and glial XBP-1s animals transgenic for VIT-2::GFP at day 1 of adulthood grown on OP50 *E. coli*. White boxes indicate eggs within the adult animal. Scale bar, 250 µM. (F) Quantification of rough ER cisternae space circularity in wild-type and glial XBP-1s animals from Figure 1J. A circularity value of 1.0 indicates a perfect circle. As the value approaches 0.0, it indicates an increasingly elongated polygon. Plots are of measurements of rough ER from n > 15 samples over 3 independent replicates. Statistics by Mann-Whitney test, p < 0.0001 (****).

**Figure S2.**
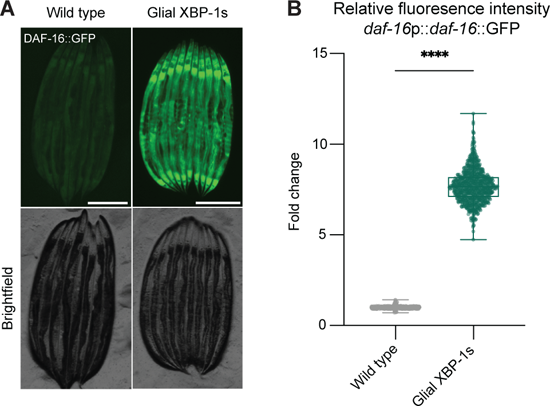
(A) Representative fluorescent micrograph of wild-type and glial XBP-1s animals transgenic for DAF-16::GFP. Animals were imaged at day 1 of adulthood. Images are representative of three independent replicates. Scale bar, 250 µM. (B) Quantification of animals in (A) using a COPAS BioSorter for whole worm fluorescence quantification. Fluorescence intensity was compared to mean of wild-type animals to determine fold change. Box plot shows median, whiskers are minimum to maximum values. Plot is representative data of three independent replicates, N > 500 animals per condition. Statistics by Kruskal-Wallis with Dunn’s multiple comparison test, p < 0.0001 (****).

**Figure S3.**
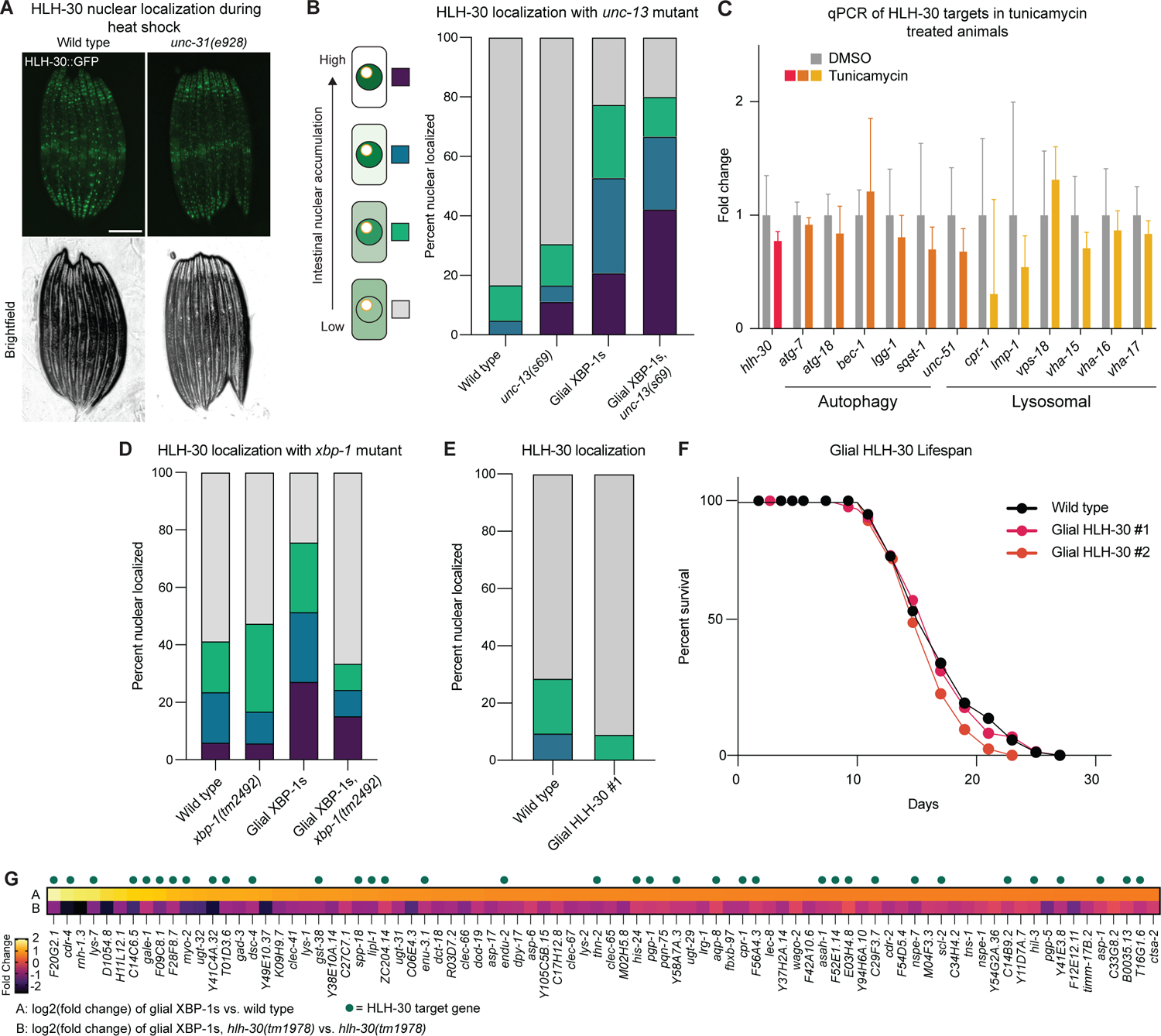
(A) Representative fluorescent micrograph of wild-type and *unc-31(e928)* mutant animals transgenic for HLH-30::GFP treated with a heat shock exposure at 35°C for 3 hours. Scale bar, 250 µM. Image is representative of 2 independent replicates. All animals imaged showed nuclear localization, N > 20 per condition. (B) Nuclear translocation of HLH-30::GFP in intestinal cells of day 2 adult wild-type and glial XBP-1s animals fed OP50 *E. coli* with and without the loss-of-function *unc-13* mutant, *unc-13(s69)*, which disrupts both isoforms of *unc-13.* N > 36 animals per condition from three independent replicates. Statistics done by Chi-squared test for independence with adjusted residuals and Bonferroni correction, statistics shown in Table S5. (C) qRT-PCR analysis of HLH-30 autophagy and lysosomal target genes from whole animal samples of wild-type animals treated with DMSO or tunicamycin. Bar graph shows mean transcript levels normalized to expression in wild-type animals treated with DMSO, from three independent biological and four technical replicates. Error bars represent SEM. Statistics by Two-way ANOVA with Sidák’s multiple comparison test, p > 0.05 (ns). All comparisons were found to be non-significant. (D) Nuclear translocation of HLH-30::GFP in intestinal cells of day 2 adult wild-type and glial XBP-1s animals fed OP50 *E. coli* with and without the loss-of-function *xbp-1* mutant, *xbp-1(tm2492)*. N > 33 animals per condition from three independent replicates. Statistics done by Chi-squared test for independence with adjusted residuals and Bonferroni correction, statistics shown in Table S5. (E) Nuclear translocation of HLH-30::GFP in intestinal cells of day 2 adult wild-type and extra chromosomal array glial HLH-30 (*hlh-17*p::*hlh-30*) fed OP50 *E. coli*. N > 30 animals per strain from two independent replicates. (F) Survival of wild-type and glial HLH-30 (*hlh-17*p::*hlh-30*, extra chromosomal arrays) animals on control RNAi at 20°C. Graph was plotted as Kaplan-Meier survival curves, and p values were calculated by Mantel-Cox log-rank test. See Table S1 for lifespan statistics. (G) Comparison of log2(fold change) of the 86 upregulated DEGs in glial XBP-1s animals compared to wild-type animals (top) compared to the log2(fold change) of these genes in glial XBP-1s animals with a loss-of-function *hlh-30* mutation, *hlh-30(tm1978)*, compared to expression in *hlh-30(tm1978)* mutants alone (bottom). log2(fold change) is color coded via a heatmap from warm (up-regulated) to cool (down-regulated) colors. Green dots above the heat map represent HLH-30 target genes. Statistics shown in Table S6.

**Figure S4.**
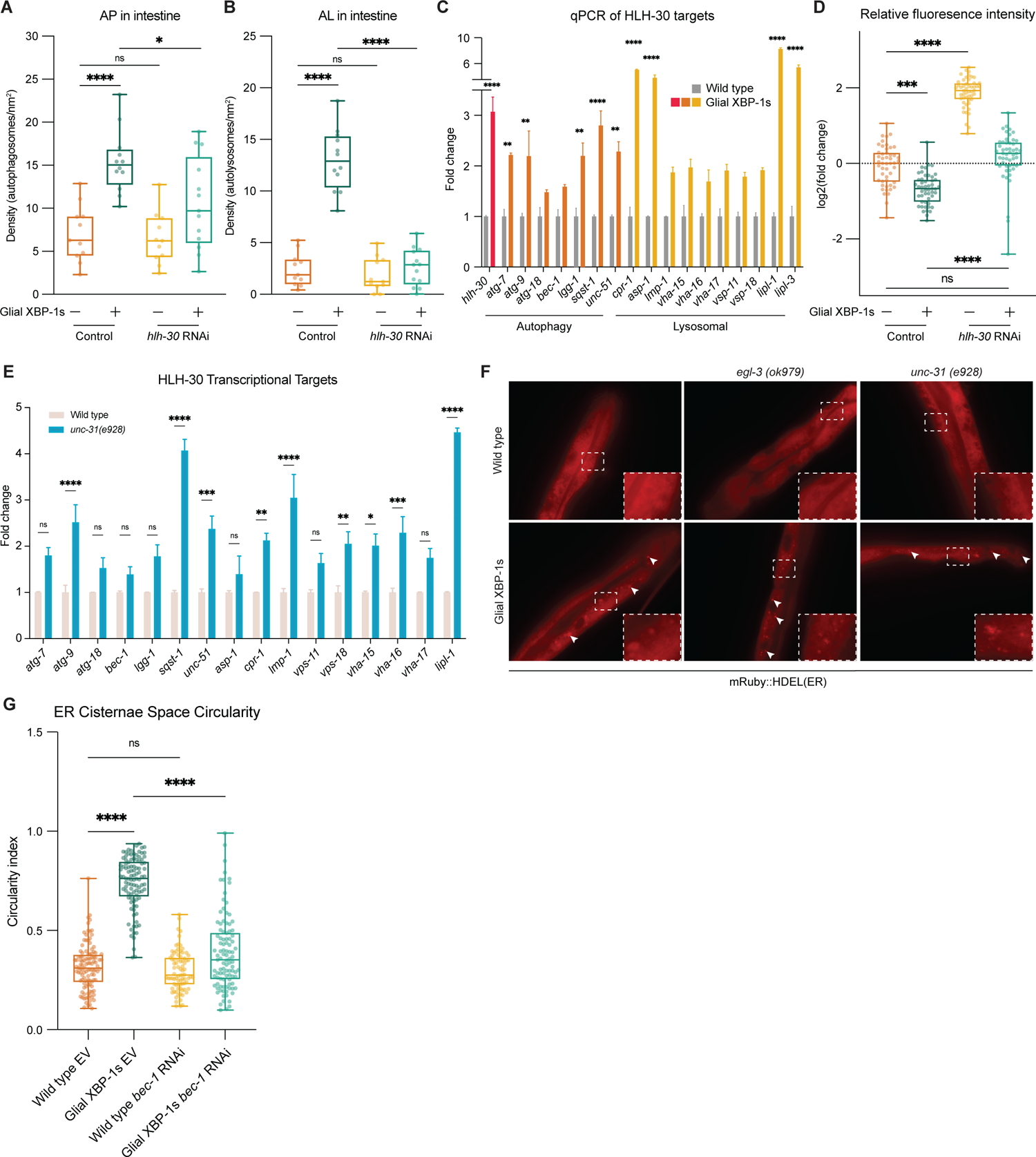
(A) Quantification of mCherry puncta co-localized with GFP (autophagosomes (AP)) intestine of wild-type and glial XBP-1s transgenic animals at day 2 of adulthood. Data are from three independent experiments, each with ≥ 5 animals. Statistics by Kruskal-Wallis with Dunn’s multiple comparison test, p < 0.0001 (****), p < 0.5 (*), p > 0.5 (ns). (B) Quantification of puncta containing mCherry alone (autolysosome (AL)) (in the intestine of wild-type and glial XBP-1s transgenic animals at day 2 of adulthood. Data are from three independent experiments, each with ≥ 5 animals. Statistics by Kruskal-Wallis with Dunn’s multiple comparison test, p < 0.0001 (****), p > 0.5 (ns). (C) qRT-PCR analysis of HLH-30 autophagy and lysosomal target genes in whole animal samples from wild-type and glial XBP-1s animals. Bar graph represents mean transcript levels normalized to wild-type control, from three independent biological and four technical replicates. Error bars represent SEM. Statistics by Two-way ANOVA with with Sidák’s multiple comparison test, p < 0.01 (**), p < 0.0001 (****). (D) Fluorescence intensity of intestinal polyQ_44_ aggregates from wild-type and glial XBP-1s animals grown on control or *hlh-30* RNAi quantified from Figure 4F using ImageJ and expressed as fluorescence intensity relative to the average intensity of wild-type animals expressing polyQ_44._ Box plot shows median, whiskers are minimum to maximum values. Statistics by Kruskal-Wallis with Dunn’s multiple comparison test p < 0.001 (***), p < 0.0001 (****). N = 50 animals per condition. (E) qRT-PCR analysis of HLH-30 autophagy and lysosomal target genes in whole animal samples from wild-type and *unc-31(e928)* animals. Bar graph represents mean transcript levels normalized to wild-type control, from three independent biological and four technical replicates. Error bars represent SEM. Statistics by Two-way ANOVA with with Sidák’s multiple comparison test, p < 0.01 (**), p <0.001 (***), p <0.0001 (****). (F) Representative micrographs from intestine of wild-type and glial XBP-1s animals transgenic for the ER marker *vha-6p*::ERss::mRuby::HDEL with or without *egl-3(ok979)* or *unc-31(e928)* mutations at day 3 of adulthood. Animals were grown on OP50 *E. coli*. ER puncta is denoted by white arrowheads. (G) Quantification of rough ER cisternae space circularity from Figure 5G in wild-type and glial XBP-1s animals fed control or *bec-1* RNAi. A circularity value of 1.0 indicates a perfect circle. As the value approaches 0.0, it indicates an increasingly elongated polygon. Plots are of measurements of rough ER from n > 15 samples over 3 independent experiments. Statistics by Kruskal-Wallis with Dunn’s multiple comparison test, p < 0.0001 (****), p > 0.5 (ns).

## SUPPLEMENTAL TABLES

**Table S1.**
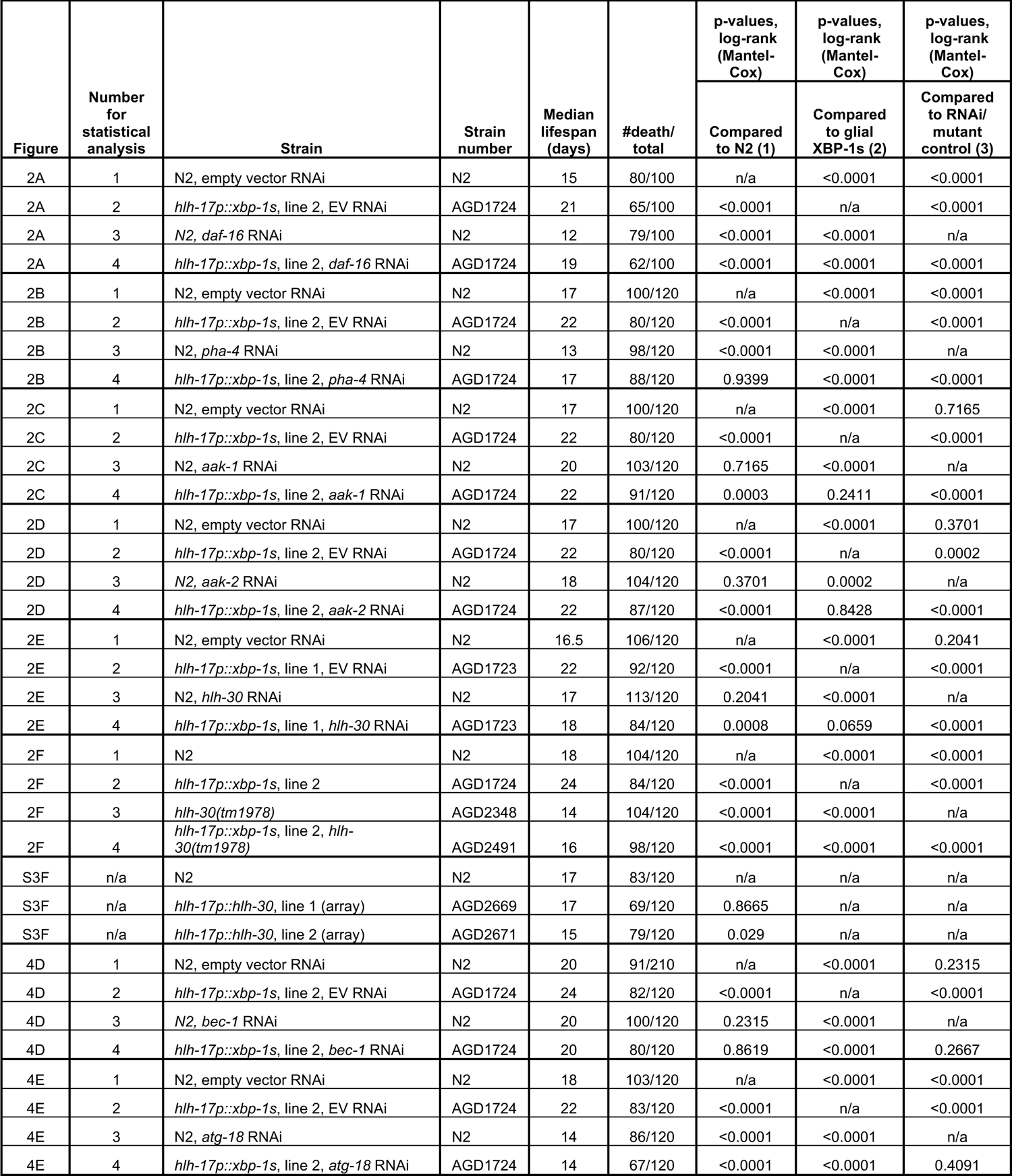
Lifespan statistics. Summary of data from representative lifespan experiments displayed in figures.

| Figure | Number for statistical analysis | Strain | Strain number | Median lifespan (days) | #death/ total | p-values, log-rank (Mantel-Cox) | p-values, log-rank (Mantel-Cox) | p-values, log-rank (Mantel-Cox) |
| --- | --- | --- | --- | --- | --- | --- | --- | --- |
|  |  |  |  |  |  | Compared to N2 (1) | Compared to glial XBP-1s (2) | Compared to RNAi/ mutant control (3) |
| 2A | 1 | N2, empty vector RNAi | N2 | 15 | 80/100 | n/a | <0.0001 | <0.0001 |
| 2A | 2 | *hlh-17p::xbp-1s*, line 2, EV RNAi | AGD1724 | 21 | 65/100 | <0.0001 | n/a | <0.0001 |
| 2A | 3 | N2, *daf-16* RNAi | N2 | 12 | 79/100 | <0.0001 | <0.0001 | n/a |
| 2A | 4 | *hlh-17p::xbp-1s*, line 2, *daf-16* RNAi | AGD1724 | 19 | 62/100 | <0.0001 | <0.0001 | <0.0001 |
| 2B | 1 | N2, empty vector RNAi | N2 | 17 | 100/120 | n/a | <0.0001 | <0.0001 |
| 2B | 2 | *hlh-17p::xbp-1s*, line 2, EV RNAi | AGD1724 | 22 | 80/120 | <0.0001 | n/a | <0.0001 |
| 2B | 3 | N2, *pha-4* RNAi | N2 | 13 | 98/120 | <0.0001 | <0.0001 | n/a |
| 2B | 4 | *hlh-17p::xbp-1s*, line 2, *pha-4* RNAi | AGD1724 | 17 | 88/120 | 0.9399 | <0.0001 | <0.0001 |
| 2C | 1 | N2, empty vector RNAi | N2 | 17 | 100/120 | n/a | <0.0001 | 0.7165 |
| 2C | 2 | *hlh-17p::xbp-1s*, line 2, EV RNAi | AGD1724 | 22 | 80/120 | <0.0001 | n/a | <0.0001 |
| 2C | 3 | N2, *aak-1* RNAi | N2 | 20 | 103/120 | 0.7165 | <0.0001 | n/a |
| 2C | 4 | *hlh-17p::xbp-1s*, line 2, *aak-1* RNAi | AGD1724 | 22 | 91/120 | 0.0003 | 0.2411 | <0.0001 |
| 2D | 1 | N2, empty vector RNAi | N2 | 17 | 100/120 | n/a | <0.0001 | 0.3701 |
| 2D | 2 | *hlh-17p::xbp-1s*, line 2, EV RNAi | AGD1724 | 22 | 80/120 | <0.0001 | n/a | 0.0002 |
| 2D | 3 | N2, *aak-2* RNAi | N2 | 18 | 104/120 | 0.3701 | 0.0002 | n/a |
| 2D | 4 | *hlh-17p::xbp-1s*, line 2, *aak-2* RNAi | AGD1724 | 22 | 87/120 | <0.0001 | 0.8428 | <0.0001 |
| 2E | 1 | N2, empty vector RNAi | N2 | 16.5 | 106/120 | n/a | <0.0001 | 0.2041 |
| 2E | 2 | *hlh-17p::xbp-1s*, line 1, EV RNAi | AGD1723 | 22 | 92/120 | <0.0001 | n/a | <0.0001 |
| 2E | 3 | N2, *hlh-30* RNAi | N2 | 17 | 113/120 | 0.2041 | <0.0001 | n/a |
| 2E | 4 | *hlh-17p::xbp-1s*, line 1, *hlh-30* RNAi | AGD1723 | 18 | 84/120 | 0.0008 | 0.0659 | <0.0001 |
| 2F | 1 | N2 | N2 | 18 | 104/120 | n/a | <0.0001 | <0.0001 |
| 2F | 2 | *hlh-17p::xbp-1s*, line 2 | AGD1724 | 24 | 84/120 | <0.0001 | n/a | <0.0001 |
| 2F | 3 | *hlh-30(tm1978)* | AGD2348 | 14 | 104/120 | <0.0001 | <0.0001 | n/a |
| 2F | 4 | *hlh-17p::xbp-1s*, line 2, *hlh-30(tm1978)* | AGD2491 | 16 | 98/120 | <0.0001 | <0.0001 | <0.0001 |
| S3F | n/a | N2 | N2 | 17 | 83/120 | n/a | n/a | n/a |
| S3F | n/a | *hlh-17p::hlh-30*, line 1 (array) | AGD2669 | 17 | 69/120 | 0.8665 | n/a | n/a |
| S3F | n/a | *hlh-17p::hlh-30*, line 2 (array) | AGD2671 | 15 | 79/120 | 0.029 | n/a | n/a |
| 4D | 1 | N2, empty vector RNAi | N2 | 20 | 91/210 | n/a | <0.0001 | 0.2315 |
| 4D | 2 | *hlh-17p::xbp-1s*, line 2, EV RNAi | AGD1724 | 24 | 82/120 | <0.0001 | n/a | <0.0001 |
| 4D | 3 | N2, *bec-1* RNAi | N2 | 20 | 100/120 | 0.2315 | <0.0001 | n/a |
| 4D | 4 | *hlh-17p::xbp-1s*, line 2, *bec-1* RNAi | AGD1724 | 20 | 80/120 | 0.8619 | <0.0001 | 0.2667 |
| 4E | 1 | N2, empty vector RNAi | N2 | 18 | 103/120 | n/a | <0.0001 | <0.0001 |
| 4E | 2 | *hlh-17p::xbp-1s*, line 2, EV RNAi | AGD1724 | 22 | 83/120 | <0.0001 | n/a | <0.0001 |
| 4E | 3 | N2, *atg-18* RNAi | N2 | 14 | 86/120 | <0.0001 | <0.0001 | n/a |
| 4E | 4 | *hlh-17p::xbp-1s*, line 2, *atg-18* RNAi | AGD1724 | 14 | 67/120 | <0.0001 | <0.0001 | 0.4091 |

**Table S2. Differentially expressed genes (DEGs) from RNAseq.** (Excel file) List of DEGs from comparison of 1) glial XBP-1s animals compared to wild type, 2) glial XBP-1s animals with *hlh-30(tm1978)* mutation compared to *hlh-30(tm1978)* alone and 3) glial XBP-1s animals with *hlh-30(tm1978)* mutation compared to glial XBP-1s animals.

**Table S3. Description of upregulated HLH-30/TFEB target genes in glial XBP-1s animals.** (Excel file) List of the 36 HLH-30/TFEB target genes upregulated in glial XBP-1s animals. Table lists gene name, GO biological process, GO cellular component, GO molecular function, and Wormbase description per gene.

**Table S4.**
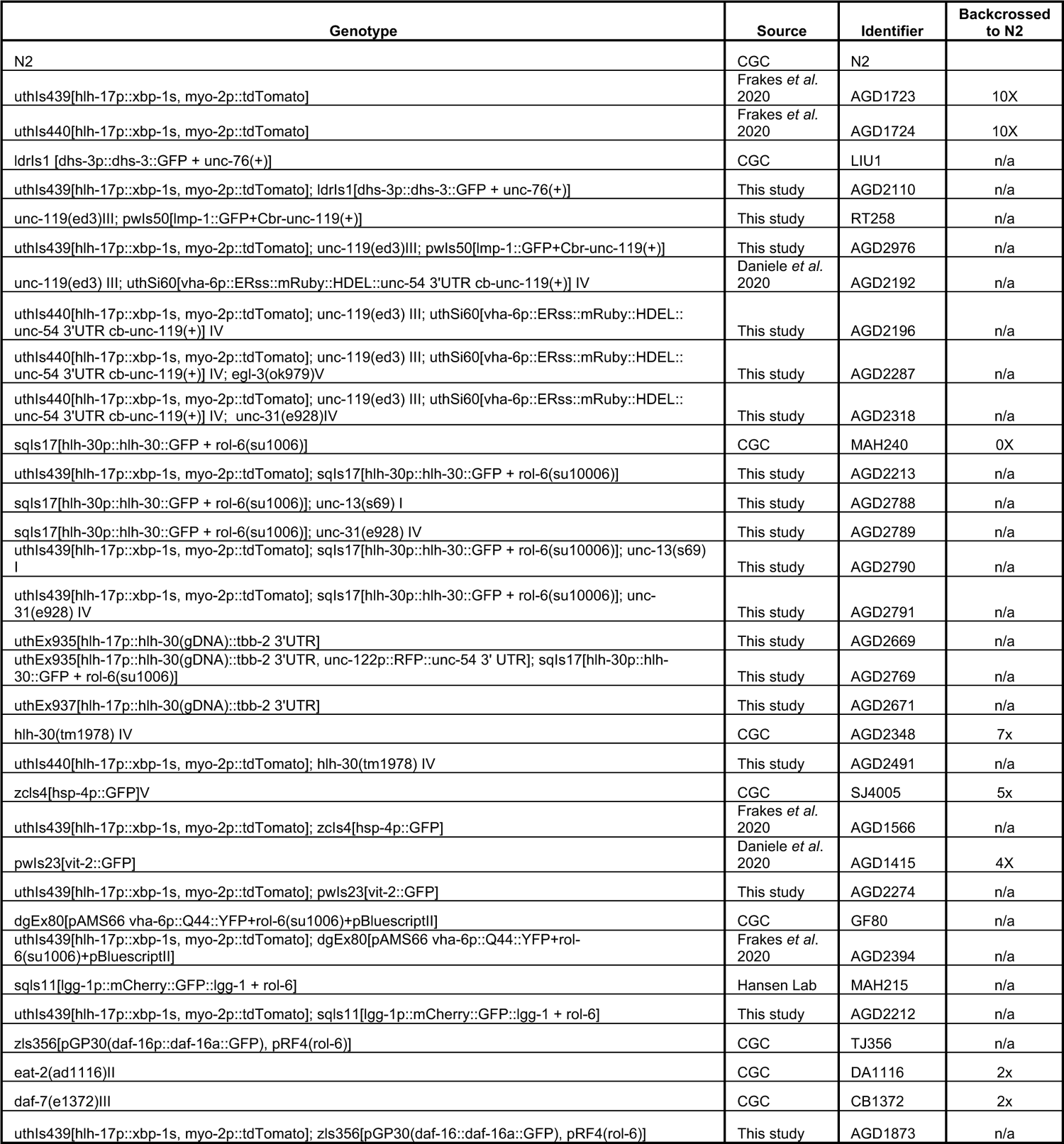
List of *C. elegans* strains used in study.

**Table S5. Statistics of stacked bar charts.** (Excel file) Data showing statistics of bar chart data (Figures 3D, S3B, S3D, and 4D). Statistics done by Chi-squared test for independence with adjusted residuals and Bonferroni correction, using the test statistic Z to test for equality of independent proportions. Values were found to be either within, statistically below, or statistically above the expected range.

**Table S6. RNAseq heat map statistics.** (Excel file) Chart showing the fold change and adjusted p values of the genes shown in the heat maps in Figures 3G and Figures S3G.

